# Lock, Protect, and Bind: In Vitro Selection of LNA-modified Aptamers Using a Mutant T7 RNA Polymerase

**DOI:** 10.1101/2025.04.09.647959

**Authors:** Kevin Neis, Laia Civit, Kathrine Pedersen, Michael Schachtner, Jesper Wengel, Ralf Wagner, Steffen Thiel, Jørgen Kjems, Julián Valero

**Affiliations:** Interdisciplinary Nanoscience Center, Aarhus University, Aarhus, Denmark; Department of Molecular Biology and Genetics, Aarhus University, Aarhus, Denmark; Department of Biomedicine, Aarhus University, Aarhus, Denmark; Institute of Medical Microbiology & Hygiene, Molecular Microbiology (Virology), University of Regensburg, Regensburg, Germany; Institute of Clinical Microbiology & Hygiene, University Hospital Regensburg, Regensburg, Germany; Biomolecular Nanoscale Engineering Center, Department of Physics, Chemistry and Pharmacy, University of Southern Denmark, Odense, Denmark

## Abstract

RNA therapeutics are powerful tools for gene modulation and targeted therapies, but their clinical application is hindered by nuclease degradation and immunogenicity. Incorporating chemical modifications, like locked nucleic acids (LNAs), can enhance nuclease resistance, targeting properties, and thermal stability. Traditionally, LNA incorporation has relied on solid-phase synthesis of short RNAs. Engineered polymerases capable of incorporating xenonucleic acids (XNAs), including LNA, into longer RNAs have been described. However, their XNA yield is limited by primer and template copy numbers, and the generated DNA-XNA duplexes can be difficult to purify.

We present a novel approach for incorporating LNA-ATP and LNA-TTP alongside 2’Fluoro (2’F)-modified pyrimidines via *in vitro* transcription using a mutant T7 RNA polymerase. This method enables efficient, primer-independent synthesis and amplification of LNA-modified RNA with low error rates.

To demonstrate its utility, we performed *in vitro* selection (SELEX) of LNA- and 2’F-modified aptamers targeting Influenza hemagglutinin (HA) and human CD40 ligand (hCD40L), two therapeutically relevant proteins. Iterative SELEX cycles yielded aptamers with low-nanomolar affinities, high specificity, and high nuclease resistance. Overall, this approach provides a scalable and versatile platform for generating chemically stabilized RNAs, fully compatible with SELEX, and holds potential for developing next-generation RNA-based therapeutics with improved pharmacokinetics.

## Introduction

RNA-based therapeutics have emerged as powerful biomedical tools, recently showcased by the great success of messenger RNA (mRNA)-based COVID-19 vaccines and their broad application in treating a panoply of diseases over recent decades. ^[1-4]^ While significant attention has been directed towards small interfering RNAs (siRNAs)^[5]^ and mRNAs^[6-9]^, there is an increasing interest in the development of aptamers. Aptamers are short single-stranded oligonucleotides (RNA or DNA) that can bind to selected targets with high affinity and specificity.^[10-12]^ They are generated via *in vitro* selection through a process termed SELEX (systematic evolution of ligands by exponential enrichment). Due to their unique binding properties, aptamers have been applied for targeting therapeutically relevant biomolecules for diagnostics and treatment.

Despite recent advances in RNA therapeutics, a major concern that greatly limits their applicability is the low chemical stability of native RNAs against nuclease degradation and their immunogenicity. To overcome these challenges, a wide variety of base and sugar modifications have been incorporated into antisense oligonucleotides (ASOs), siRNAs and mRNAs.^[7, 13]^ Locked nucleic acids (LNAs) stand among the most notable nucleotide modifications in oligonucleotides and are extensively used in therapeutic ASOs. LNAs feature a bridge, commonly a methylene linker, between the 2’-O and the 4’-C positions of the ribose ring, “locking” the sugar in the 3’-endo conformation.^[14-16]^ This prevents nuclease degradation and confers great base-pairing stability and enhanced targeting, with a substantial increase in melting temperature (Tm) of the duplex.^[17-19]^

Until recently, the efficient incorporation of LNA modifications into RNA was predominantly limited to the production of relatively short oligonucleotides via complex solid-phase synthesis.^[18]^ This restricts the generation of longer, functional LNA-modified oligonucleotides or the production of aptamers, which require polymerases capable of synthesizing and reading LNA-modified sequences during iterative cycles of in vitro selection. To date, only two engineered polymerases have been reported to generate LNA-modified oligonucleotides via primer extension reaction (PER). ^[20-21]^ These enzymes efficiently produce fully modified LNA-oligonucleotides and allow combining LNA and 2’-O-methyl (2’-OMe) modifications with relatively high fidelity.^[22]^ Hoshino *et al*. demonstrated that engineered *thermococcus kodakaraensis* (KOD) DNA polymerases capable of LNA incorporation could be employed for aptamer selection and reported an LNA-modified aptamer binding with nanomolar affinity against thrombin using capillary electrophoresis.^[21]^ While remarkable, RNA amounts produced by PER reactions are inherently limited by primer and template concentrations, unlike RNA polymerases, which produce multiple RNA copies from a single template. Additionally, PER requires labor-intensive sample processing and purification to remove the template from the reaction.^[23]^ Moreover, some engineered polymerases have shown slow kinetics, reduced processivity and fidelity issues compared to their wild-type analogs.

Alternatively, enzymatic LNA incorporation into RNA has been explored using terminal deoxynucleotidyl transferases or template-independent poly(U) polymerases.^[24]^ However, these polymerases are impractical for the synthesis of LNA modified oligonucleotides, since the N+1 incorporation efficiency rarely exceeds 60-70%. Veduu *et al*. reported the enzymatic incorporation of LNA modifications into oligonucleotides through PCR amplification using DNA polymerases and co-transcriptional incorporation into RNA using wild-type T7 RNA polymerase (T7RNAP).^[25]^ Despite these advancements, the polymerases exhibited moderate efficiencies and generated only short oligonucleotides with limited amount of LNA modifications.

Here, we report the generation of LNA-modified oligonucleotides via transcription using a mutant T7RNAP (Y639F). ^[26]^ This approach enables the incorporation of LNA adenosine (LNA-A) and LNA thymidine (LNA-T), in combination with 2’-fluoro (2’F) pyrimidine modifications, with high efficiency and fidelity, while remaining fully compatible with *in vitro* evolution techniques. To showcase the broad applicability of our method, we demonstrate the selection of LNA-modified aptamers that bind with high affinity to two biologically relevant protein targets: hemagglutinin, a surface glycoprotein of the Influenza virus,^[27]^ and CD40 ligand (CD40L, also known as CD154), a key immunomodulatory protein expressed by a subset of T -cells, essential for B-cell activation and differentiation.^[28-29]^ The selected LNA and 2’F-modified aptamers exhibit nanomolar affinities to these structurally and functionally diverse targets, along with superior stability against nuclease degradation.

## Results and Discussion

### Assessing LNA Incorporation Efficiency and Fidelity in RNA Transcripts

To assess the feasibility of generating LNA- and 2’F-modified RNA for aptamer selection using the Y639F T7RNAP mutant, we employed two DNA template libraries “N40-mA” and “N40-mU”. These were designed to accommodate *in vitro* transcription (IVT) and subsequent reverse transcription (RT) – two crucial steps in SELEX. The design entails a 40-nucleotide random region flanked by constant regions, devoid of adenosines or thymidines, respectively (Figure 1A, Table S1). This exclusion of LNA modifications aims to reduce excessive template-primer stability during reverse transcription, mitigate sequence bias in LNA incorporation and prevent complications during transcription initiation.

**Figure 1.**
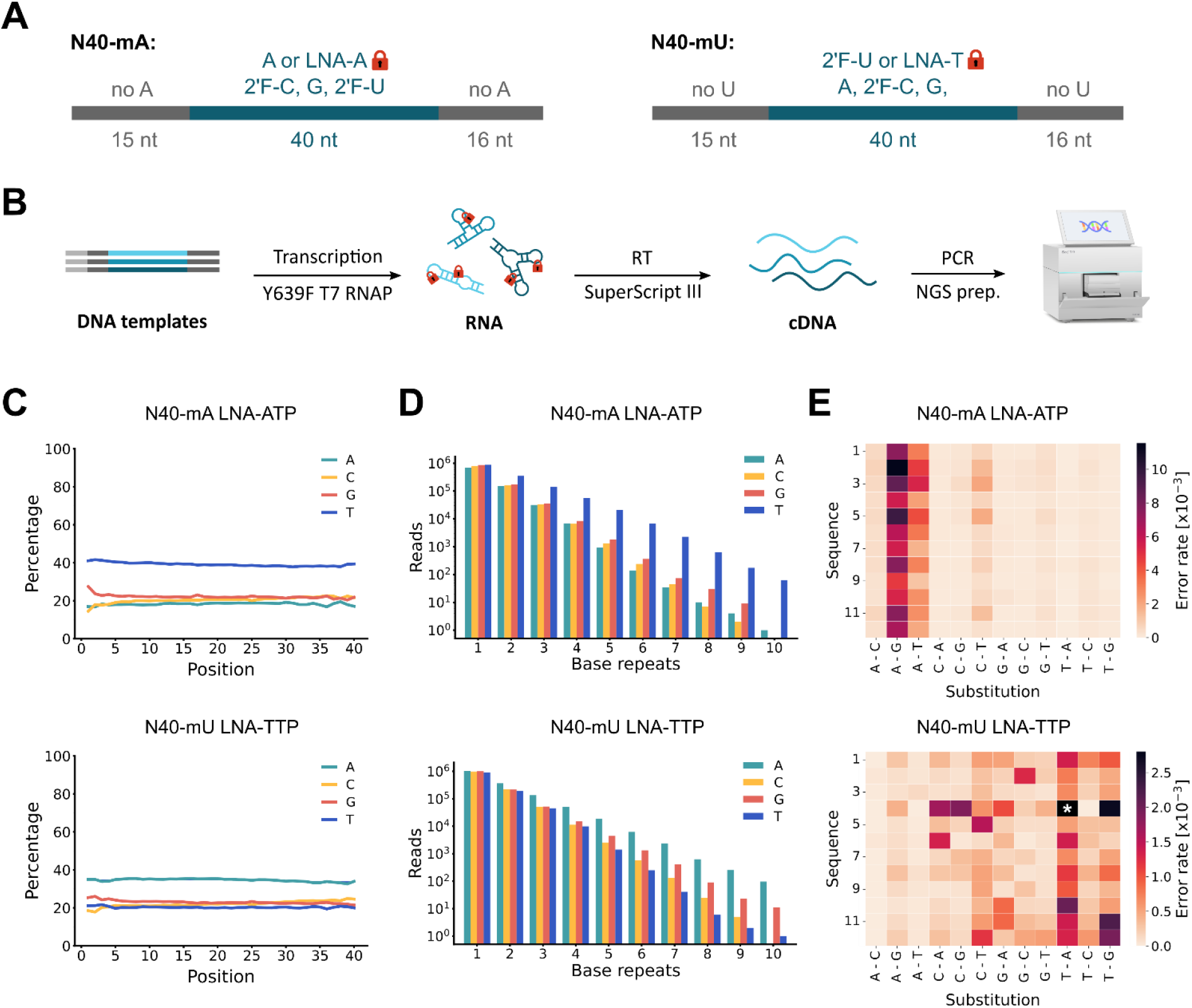
NGS analysis of LNA-ATP and LNA-TTP incorporation alongside 2’F-pyrimidines into RNA. (A) Schematic representation of the N40-mA and N40-mU RNA libraries. Constant regions are shown in grey, while the random regions and incorporated nucleotides are highlighted in cyan. nt = nucleotide. (B) Workflow for the NGS analysis, including *in vitro* transcription (IVT), reverse transcription (RT), polymerase chain reaction (PCR), and NGS library preparation. LNA-modified nucleotides are illustrated by red locks. (C) Nucleotide distribution in the “N40-mA” (LNA-ATP) and “N40-mU” (LNA-TTP) DNA libraries after one round of IVT, RT and PCR across the 40-nt random region. (D) Abundance of the indicated number of consecutive base repeats in the sequenced DNA libraries. (E) Error rates of LNA-ATP and LNA-TTP incorporation after one round of IVT, RT and PCR for each of the twelve selected sequences (Table S1). Error rates for each substitution (template base – sequenced base) are shown relative to the total number of nucleotides analysed per sequence. (*) For N40-mU (LNA-TTP), the scale is limited to 2.8*10^−3^, but the T-A substitution for sequence 4 has an error rate of 7.7 *10^−3^.

First, we examined the IVT yield as a measure of LNA incorporation efficiency. IVT of N40-mA using LNA-ATP, 2’F-pyrimidines and N40-mU using LNA-TTP and 2’F-cytidines, yielded 31% and 49% full-length RNA, respectively, compared to transcriptions containing only 2’F-pyrimidines, which are known to be efficiently incorporated by Y639F T7RNAP^[26]^ (Figure S1). Incorporation of LNA-CTP, LNA-GTP, or a combination of LNA-ATP and LNA-TTP resulted in either low or no transcription yields for both libraries. Furthermore, incorporating LNA-ATP into N40-mU and LNA-TTP into N40-mA proved inefficient and yielded shorter RNA fragments, suggesting that LNA modifications in the constant regions reduce transcription efficiency.

Subsequently, we employed next-generation sequencing (NGS) to provide detailed insights into potential biases and misincorporation of LNA-NTPs during transcription and reverse transcription (RT). To this end, RNA was reverse transcribed using SuperScript III,^[30]^ cDNA was amplified by PCR using Phusion polymerase, and the amplicons were analyzed by NGS (Figure 1B).

LNA incorporation introduces a shift in nucleotide frequency (Figure 1C). While the initial DNA pools and transcripts lacking LNA-NTP modifications display a nearly equal nucleotide distribution across the random region (Figure S2A), the introduction of LNA-ATP in the N40-mA library increases the thymidine content by approximately 14% (Figure 1C). Similarly, LNA-TTP incorporation in the N40-mU library raises the adenosine content by around 12%. The remaining nucleotides, including LNA-modified ones, showed a comparable distribution, with a slight decrease in (2-5%) relative to non-LNA transcripts (Figure 1C). Notably, no positional bias in nucleotide distribution was observed across the sequence length. Additionally, we evaluated the incorporation efficiency of LNA-CTP into the N40-mU library, which resulted in a substantial 15% decline in cytidine frequency (Figure S2C). Consequently, further investigations of LNA-CTP were discontinued.

We further analyzed the prevalence of consecutive LNA-nucleotide incorporations, as the efficiency of incorporating successive LNA modifications might progressively be reduced (Figures 1D and S2B). As expected, we observed a slight decrease in the number of sequences carrying a high number of consecutive LNA modifications, consistent with the shifted nucleotide distribution. However, the sequence diversity in the libraries remained high (Table S2), with some sequences containing up to ten consecutive LNA modifications (Figure 1D). Moreover, the observed increase in consecutive T and A motifs for RNA modified with LNA-ATP and LNA-TTP, respectively, correlated well with the higher T or A content in the libraries.

To assess the combined fidelity of the T7 RNAP mutant in incorporating LNA-modified nucleotides and the accuracy of SuperScript III in reading LNA-modified RNA, we evaluated the cumulative error rates of transcription, RT and PCR using NGS. For this, we used twelve templates with known sequences, flanked by the constant regions of N40-mA and N40-mU for LNA-modified As and Ts, respectively (Table S1). These sequences were randomly generated, ensuring equal nucleotide ratios, with each possible base triplet represented at least once. To confirm the absence of residual DNA templates, we performed direct PCR amplification of the purified RNA samples prior to the RT step. No PCR product was observed in these samples, while robust amplification was detected in the reverse-transcribed samples, confirming both the absence of residual DNA and the requirement of cDNA for amplification (Figure S3).

Error rate analysis for each sequence revealed that A→G and A→T substitutions were the most prevalent when incorporating LNA-ATP alongside 2’F-C and 2’F-U in the twelve N40-mA sequences, with accumulated error rates ranging from 1.2 × 10^−3^ to 11.5 × 10^−3^ (Figure 1E). This indicates a reliable LNA-ATP incorporation. For LNA-TTP incorporation in combination with 2’F-C in the twelve N40-mU sequences, we observed considerably lower error rates, with T→A and T→G substitutions being the most common. However, one sequence exhibited higher error rates, likely due to transcription or RT bias, as it had significantly fewer reads than compared to the control pool (Table S3). The low error rates found for the 2’F-modified controls (Figure S4A) suggest that the observed errors primarily stem from mistakes in LNA incorporation. While we predominantly observed substitutions of LNA-modified bases to purines, both LNA-modified pools showed a significant number of C→T mutations, absent in the 2’F-modified control transcriptions (Figure S4B), which is especially surprising for the LNA-T modified sample.

Our results demonstrate that commercially available enzymes, such as the Y639F T7RNAP mutant and SuperScript III reverse transcriptase, efficiently enable the combination of LNA and 2’F chemistries to generate and read RNAs, respectively, with low overall error rates - 11.7 × 10^−3^ for LNA-ATP and 4.4 × 10^−3^ for LNA-TTP (Figure S4C and Table S4). These error rates are significantly lower than those reported for engineered polymerases that enable combined incorporation of LNA and 2’-OMe modifications via PER^[20-22]^. Furthermore, the overall accumulated error rate for 2’F-pyrimidine incorporation (0.8 × 10^−3^) is 20-27 times lower than that observed for 2’OMe-modified RNA generated by engineered polymerases^[22]^. While direct comparisons with previously reported PER systems are challenging due to differences in modification type (2’F vs. 2’OMe) and degree of modification, our data highlight the versatility of the Y639F T7RNAP mutant for simultaneous incorporation of LNA-modified A and T nucleotides alongside 2’F-pyrimidine modifications.

### In Vitro Selection of RNA Aptamers Containing LNA and 2’F-Modifications

To determine the feasibility of our method for selecting LNA- and 2’F-modified RNA aptamers, we performed SELEX against two therapeutically relevant protein targets: the Influenza virus surface protein, hemagglutinin (HA), and human CD40 ligand (hCD40L). HA mediates viral attachment to host cells, a critical step in Influenza infection.^[27]^ hCD40L, expressed on activated T cells, plays a key role in B-cell maturation and function by engaging the CD40 protein, constituting an important molecular checkpoint for immunomodulation and cell-cell communication.^[28-29]^ Based on our previous screening results, we selected RNA libraries modified with LNA-T and 2’F-C. This combination provided the highest transcription efficiency, low error rates and minimal nucleotide bias which are crucial factors for SELEX success. The SELEX workflow is illustrated in Figure 2A. Nickel-nitrilotriacetic acid (NTA)-magnetic nanoparticles were used for His-tag-based target immobilization. To remove nonspecific His-tag binders, the LNA-T- and 2’F-C-modified RNA library was first incubated with an unrelated His-tagged nanobody. The supernatant, containing unbound sequences, was subsequently incubated with NTA-magnetic nanoparticles loaded with either the T4-foldon stabilized Influenza A H1N1 HA (A/England/195/09) trimer or the trimeric hCD40L (Figure 2B). Target-bound sequences were subjected to RT-PCR amplification and subsequently transcribed to generate the library for the next selection round. The selection pressure was gradually increased by reducing the target protein concentration and enhancing washing stringency (Experimental Section, SI).

**Figure 2.**
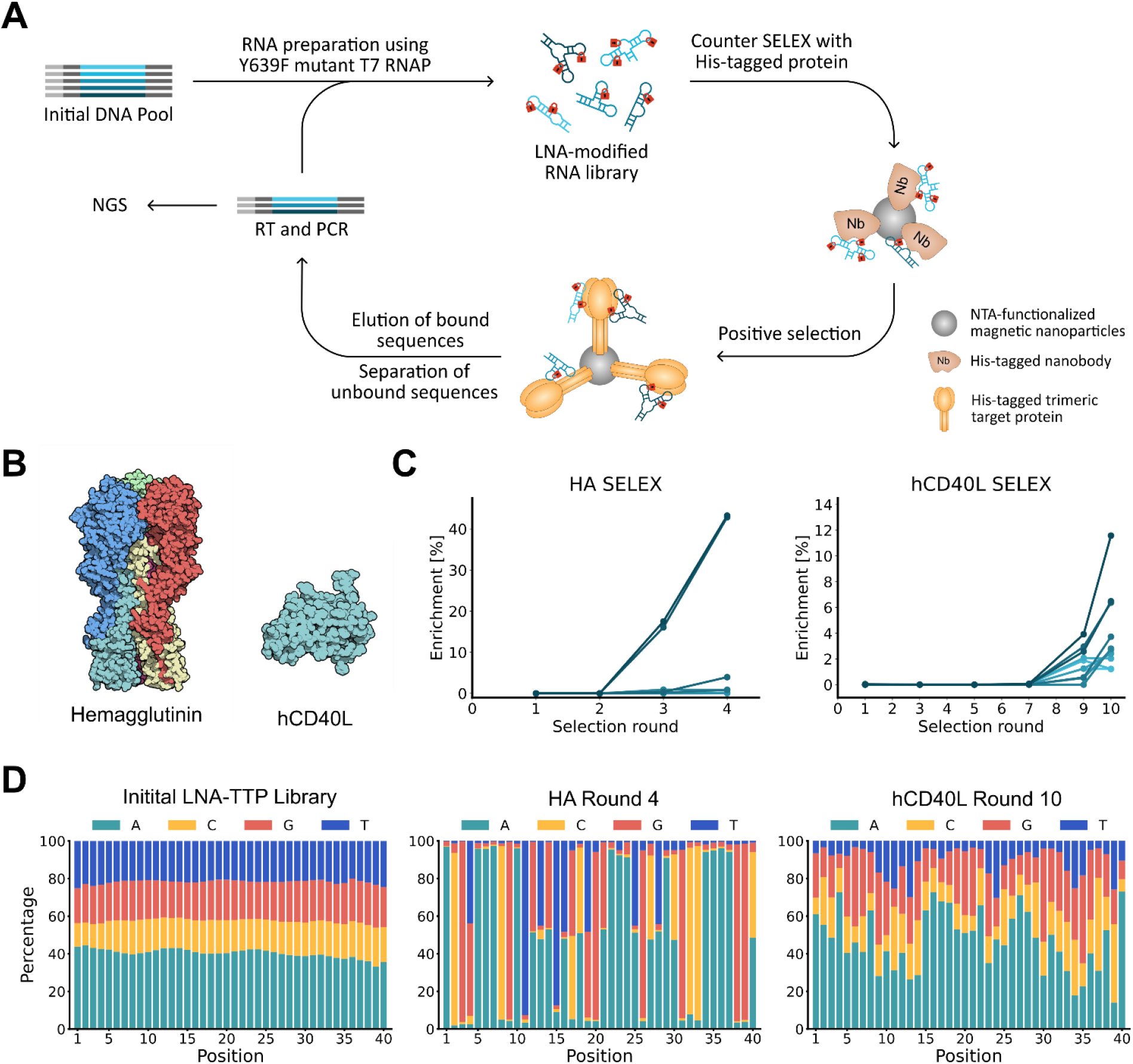
SELEX using an LNA-TTP and 2’F-CTP modified RNA library targeting Influenza hemagglutinin (HA) or human CD40L (hCD40L). (A) Schematic overview of the aptamer selection process, including a counter SELEX step using a His-tagged nanobody (Nb) to eliminate His-tag-binding RNAs. LNA-modified nucleotides are illustrated by red locks. (B) 3D structures of the target proteins: Influenza A H1N1 HA trimer [PDB: 1RUZ] and monomeric hCD40L [PDB: 1ALY]. (C) Enrichment of the ten most abundant sequences after round 4 of HA SELEX and round 10 of hCD40L SELEX, shown in comparison to each sequenced selection pool. (D) Nucleotide distribution within the 40-nt random region of the initial LNA-SELEX library and after the last SELEX round for each target. The nucleotide distributions for intermediate SELEX rounds are shown in Figure S5. The initial LNA-SELEX library was derived from a DNA template synthesized by Integrated DNA Technologies (IDT), which contained a lower overall cytidine content and a higher adenosine frequency relative to the other bases.

NGS analysis was performed after round 4 for the HA-targeted selection and round 10 for the hCD40L-targeted selection to monitor the evolution of the SELEX libraries. For the HA selection, a strong enrichment was observed, with two LNA-containing sequences dominating the library by round 3 (Figure 2C). In contrast, the enrichment for hCD40L-binding clones progressed more gradually, becoming significant only after round 10, where the most enriched sequence constituted 12% of the final library. The pronounced shift in nucleotide distribution between the initial library and the final selection rounds underscores the effective enrichment of single sequences (Figure 2D and S5). Notably, after ten selection rounds, the overall T content remained at 10% (Figure S6), confirming that LNA-T is well tolerated in the SELEX process, despite its lower representation in the initial pool.

### Binding Evaluation of Selected Aptamers to HA and hCD40L

Based on sequence enrichment, homology, and conserved sequence motifs identified in our sequencing data, we selected seven LNA- and 2’F-modified clones and produced these via IVT (Table S5). Biolayer interferometry (BLI) was used for initial screening to evaluate the binding of individual clones to their respective protein targets (Figure S7A). Among them, two clones from the fourth selection round against HA, HA-LNA-1 and HA-LNA-2, and two clones from the tenth selection round against hCD40L, hCD40L-1 and hCD40L-4, showed positive binding to their respective target proteins (Sequences in Figure 3A).

**Figure 3.**
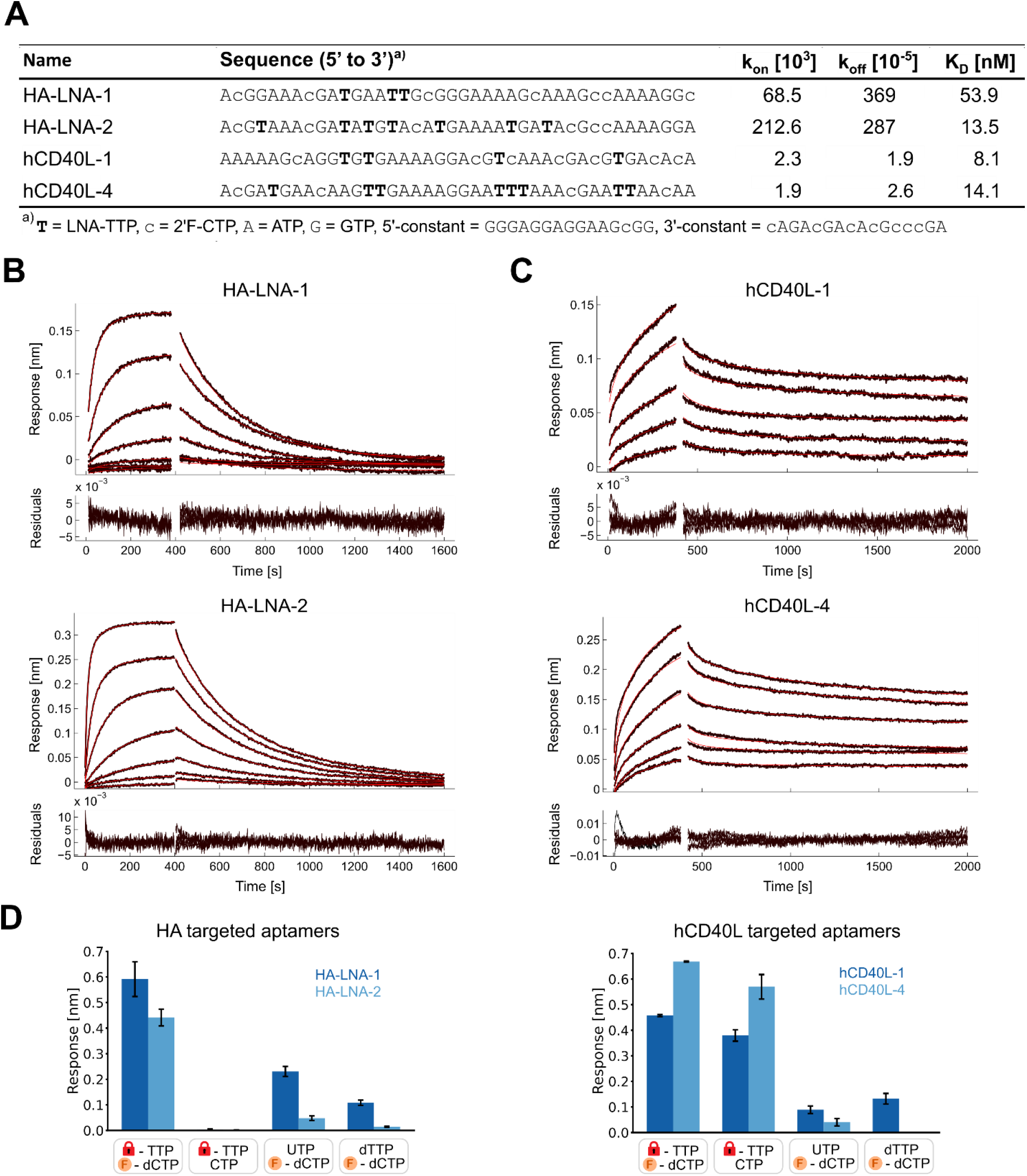
Binding studies of aptamers targeting hemagglutinin (HA) and human CD40L (hCD40L). (A) Sequences of selected aptamers targeting HA (HA-LNA-1 and HA-LNA-2) and hCD40L (hCD40L-1 and hCD40L-4), along with kinetic parameters (k_on_ and k_off_) and affinity constants (K_D_) obtained from biolayer interferometry (BLI) analysis using Evilfit[17]. Kinetic data for the most abundant species of hCD40L aptamer interactions are mentioned (Figure S8). LNA-TTP modifications are represented in bold uppercase letters, and 2’F-CTP modifications in lowercase letters. (B) BLI sensorgrams for HA-LNA-1 and HA-LNA-2 immobilized on streptavidin-coated sensors, with HA titrated in a 3-fold dilution series from 300 nM to 0.4 nM. (C) BLI sensorgrams for His-tagged hCD40L immobilized on Ni-NTA sensors, with hCD40L-1 and hCD40L-4 titrated in a 2-fold dilution series from 62.5 nM to 3.9 nM and 125 nM to 3.9 nM, respectively. (C and D) Association phase (0–400 s) and dissociation phase (400–1600/2000 s) were fitted (red lines) using Evilfit,^[31-32]^ accounting for surface effects. Residuals are displayed below. (D) Comparison of binding affinities for LNA-TTP- and 2’F-C-modified aptamers versus their nucleotide-switched variants at 500 mM HA concentration (biotinylated RNA immobilized on streptavidin sensor) or at 500 nM aptamer concentration (hCD40L immobilized on Ni-NTA sensors). BLI response after 400 s association is shown (HA: n = 4, hCD40L: n = 3). LNA-modified nucleotides are represented by a red lock, and 2’F-modified nucleotides by an orange ‘F’ symbol. Error bars represent the standard error.

To determine their binding affinities, we conducted BLI titration assays. Biotinylated HA-LNA-1 and HA-LNA-2 were immobilized on streptavidin-functionalized sensors and exposed to increasing HA concentrations. Similarly, his-tagged hCD40L was immobilized on Ni-NTA sensors and exposed to varying concentrations of hCD40L-1 and hCD40L-4. All four aptamers displayed strong and specific affinities in the low nanomolar range (Figure 3A-C, S7B). As shown in the sensorgrams (Figure 3B), the three-fold difference in affinity between HA-LNA-1 and HA-LNA-2 primarily arises from differences in association kinetics (k_on_), whereas dissociation rates (k_off_) remain comparable.

Interestingly, some sequences enriched in the final selection rounds contained a high number of LNA-T modifications. For instance, HA-LNA-2 and hCD40L-4 carried seven and eight LNA-Ts, respectively, indicating that highly LNA-modified sequences are well tolerated and can survive the selection pressure during SELEX.

To assess the impact of the LNA and 2’F-modifications on the structure and function of the selected aptamers, we investigated the effects of substituting LNA-TTP with unmodified UTP or dTTP and substituting 2’F-dCTP with unmodified CTP. For the HA aptamers, a strong decrease in binding was observed when substituting LNA-TTP with UTP or dTTP (Figure 3D). A similar effect was observed for hCD40L aptamers upon replacement of the LNA-T with unmodified U or dT, indicating that LNA modifications might play a crucial role in either stabilizing aptamer structure or mediating key interactions with the target protein. Interestingly, while the removal of the 2’F-C modification did not greatly affect hCD40L binding, it significantly lowered the binding affinity of HA aptamers. This suggests that the hydroxyl group in CTP may induce an alternative folding that disrupts HA aptamer-target interactions.

### Serum Stability Studies of Selected LNA-and 2’F-Modified Aptamers

We investigated the influence of LNA modifications on the physicochemical properties of the selected aptamers. LNA modifications are reported to enhance thermal stability and nuclease resistance of oligonucleotides.^[14, 17, 19]^ These features are crucial for their application in diagnostics using biological samples or for their potential therapeutic use. First, we assessed both the independent and synergistic effect of LNA and 2’F-modifications on the stability of the four selected aptamers against nuclease degradation, by incubating them at 37°C in 25% human serum and analyzing their degradation kinetics via gel electrophoresis (Figure 4). Aptamers containing both LNA-T and 2’F-C modifications exhibited significantly enhanced nuclease resistance, with minimal degradation (<20%) after four days. In contrast, aptamers lacking either LNA-T or 2’F-C were rapidly degraded within minutes. Aptamers containing only LNA modifications were degraded into shorter fragments (Figure S9, S10), likely due to the susceptibility of the unmodified flanking regions to exonuclease activity. Conversely, those containing only 2’F-C modifications remained more intact, producing longer degradation fragments due to the higher 2’F-C content in the flanking constant regions (Figure S9, S10).

**Figure 4.**
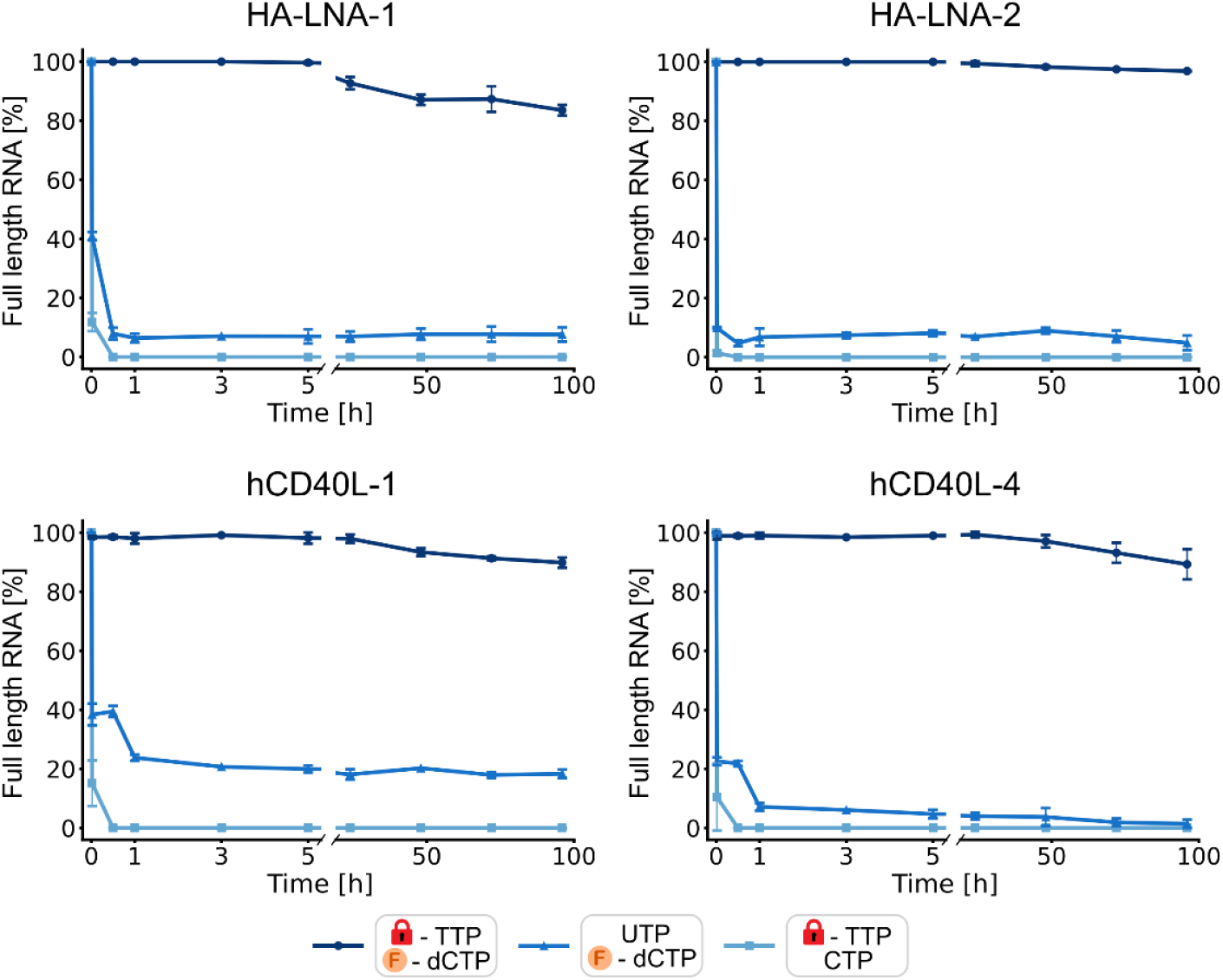
Nuclease resistance of LNA- and 2’F-modified RNA aptamers. The percentage of remaining full-length RNA for selected aptamers carrying different chemical modifications after incubation in 25% human plasma over 4 days is shown. The data were quantified from denaturing gel electrophoresis (Figures S9 and S10). LNA-modified nucleotides are represented by a red lock, and 2’F-modified nucleotides by an orange ‘F’ symbol. Error bars represent the standard error (n = 2).

To assess the effect of LNA chemistry on thermal stability, we recorded melting curves of the aptamers generated with or without LNA-T modifications by monitoring the absorbance at 260 nm across different temperatures. Although exact melting temperatures could not be accurately determined due to the complex folding profiles of these aptamers, LNA-modified HA-LNA-2 and hCD40L-4, which contain 7 and 8 LNA-Ts, respectively, showed a significant increase in thermal stability (Figure S11). In contrast, HA-LNA-1 and hCD40L-1, with fewer LNA modifications, showed less pronounced stabilization. Notably, some LNA nucleotides may reside in loops or other non-base-paired regions, contributing to aptamer structure without directly affecting thermal stability.

Conducting future SELEX experiments at elevated temperatures or within biological fluids may enhance the enrichment of aptamers, where LNA modifications significantly contribute to improved thermal stability and nuclease resistance.

## Conclusion

In this study, we demonstrated that LNA-A and LNA-T modifications can be efficiently and reliably incorporated into RNA alongside 2’-F modifications using the commercially available mutant Y639F T7RNAP. LNA-ATP incorporation and reverse transcription exhibited overall error rates comparable to those reported for other engineered polymerases, while LNA-TTP incorporation resulted in threefold lower error rates. Our method overcomes the primer and template concentration limitations of PER-based systems, enabling the amplification of LNA-containing oligonucleotides. Importantly, this approach is fully compatible with in vitro evolution techniques (SELEX), facilitating the selection of LNA- and 2’F-modified RNA aptamers with strong and specific binding in the low nanomolar range, as demonstrated for HA and hCD40L.

Since post-selection modifications can disrupt aptamer function, incorporating LNA modifications during SELEX is crucial. Our findings show that substituting LNA nucleotides with other nucleotide analogs in selected aptamers significantly reduced their binding affinity, highlighting their essential role in proper folding and target recognition. Moreover, LNA-modified aptamers show, in a sequence-dependent manner, enhanced thermal stability and protection against nuclease degradation, influenced by the number and position of LNA and 2’F-modifications.

Overall, our method provides a robust strategy for generating relatively long LNA-containing RNA transcripts while maintaining compatibility with 2’F modifications at other nucleobases. It overcomes limitations of chemical synthesis, restricted to generating shorter LNA-containing oligonucleotides, by offering a competitive approach to produce longer LNA-enhanced RNA molecules, with potential applications in gene regulation and functional targeting. Additionally, it enables the development of high-affinity, chemically stable aptamers targeting biomolecules of therapeutic interest. This opens new avenues for developing more structurally robust and stable aptamers that can potentially exhibit reproducible and consistent performance under various conditions, including elevated temperatures and harsh biological environments.

## Supporting Information

Supporting Information is available containing the Experimental Section and supplementary Figures and Tables that support the main discussion of the article.

## Acknowledgements

Lundbeck experiment grant (Grant number: R346-2020-1999), General-purpose virus neutralizing engulfing shells with modular target-specificity (VIROFIGHT) (European Union H2020-FETOPEN Grant 899619), and CEMBID (Center for Multifunctional Biomolecular Drug Design, Grant number: NNF17OC0028070). We thank Maria Gockert for assisting in the manuscript preparation. We thank Kristian Juul-Madsen for assisting with the Evilfit analysis.

