## Supplementary Information for "Lock, Protect, and Bind: In Vitro Selection of LNA-modified Aptamers Using a Mutant T7 RNA Polymerase"

### Experimental Section

All oligonucleotides, including primers and single-stranded DNA library, were synthesized by Integrated DNA Technologies (Table S6).

**Double-stranded N40-mA and N40-mU DNA library preparation by Klenow extension:** The double-stranded DNA (dsDNA) transcription templates were generated via Klenow extension. The N40-mA or N40-mU forward primer was annealed to the single-stranded N40-mU or N40-mA library, respectively, using a 1.2x molar excess (95 °C for 5 min, 50 °C for 5 min, then held at 37 °C with a ramping speed of 0.1 °C/s). Extension was performed using Klenow Fragment, exo (Thermo scientific, cat. no. EP0421) for 1 hour at 37 °C. The resulting dsDNA library was purified via 6% native poly-acrylamide gel electrophoresis, followed by passive elution in 0.3 M sodium acetate (pH 5.2) and ethanol precipitation.

**Generation of dsDNA templates by PCR:** dsDNA templates for individual selected sequences, N40-mU and N40-mA 12-clone pools, and cDNA from the SELEX process were generated via PCR. Reactions were performed using Phusion High-Fidelity DNA Polymerase (Thermo Scientific, cat. no. F530XL) with 1 µM of the respective N40-mU/N40-mA forward and reverse primers. The thermal cycling program was as follows: Initial denaturation at 95 °C for 2 min, followed by cycles of 95 °C for 30 sec, 55 °C for 30 sec, and 72 °C for 20 sec with a final extension step at 72 °C for 2 min). PCR products were purified using the GeneJET PCR purification kit (Thermo Scientific, cat. no. K0702) according to the manufacturer's instructions.

***In vitro* transcription (IVT) protocol:** All RNA libraries, 12-clone RNA pools, and individual RNA aptamers used in this study were generated by IVT from the corresponding purified dsDNA template. The transcription mix contained mutant Y639F T7 RNA polymerase (Aptamist, Denmark) in T7 transcription buffer (80 mM Hepes [pH 7.5], 25 mM $MgCl_2$, 2 mM spermidine-HCl), along with ATP and GTP (Jena Bioscience), 2'-Fluoro CTP and 2'-Fluoro UTP (Metkinin Chemistry) at 2.5 mM each. For LNA containing libraries and aptamers, ATP and 2'-Fluoro UTP were replaced by LNA-ATP or LNA-TTP, respectively, at 1.25 mM. The reaction was supplemented with 12.5 mM dithiothreitol (DTT, Invitrogen), 10% DMSO, 0.005 U/µL inorganic pyrophosphatase (IPP, Thermo Scientific), and 0.05 mg/mL bovine serum albumin (BSA), then incubated overnight at 37 °C. Following transcription, the DNA template was digested by incubation with DNase I (Thermo Scientific) at 37 °C for 20 min. RNA products were either purified by an 8% denaturing polyacrylamide gel electrophoresis, followed by passive elution and ethanol precipitation, or vie column purification using the RNA Clean & Concentrator-25 kit (Zymo Research, cat. no. R1018).

**Incorporation analysis of LNA-NTPs into RNA by mutant T7 polymerase:** The incorporation efficiency of LNA-NTPs in combination with 2'Fluoro-modified NTPs was evaluated using the N40-mU and N40-mA libraries. To determine error rates from the LNA-NTP incorporation, 12 clones with known sequences in the primer flanked regions were used (Table S1). The IVTs were performed as described above. Control samples contained ATP, GTP, 2'Fluoro-CTP and 2'Fluoro-UTP, while in LNA-modified samples 2'Fluro-UTP was replaced by LNA-TTP (N40-mU library), and ATP was replaced by LNA-ATP (N40-mA library). Following DNase I treatment and purification by denaturing poly-acrylamide electrophoresis, 25 pmol of RNA was reverse transcribed using the corresponding JL40 N40-mA/N40-mU reverse primers and SuperScript III reverse transcriptase (Invitrogen, cat.no. 18080093). cDNA was then amplified by PCR as described above. IVT yields were assessed from denaturing poly-acrylamide electrophoresis using ImageJ (https://imagej.nih.gov/ij/) and compared to the 2'F-modified control samples.

#### Generation of soluble recombinant hemagglutinin for aptamer selection

The hemagglutinin (HA) gene of A/England/195/2009 was truncated after Ala515 (mature HA numbering) to remove the transmembrane and cytoplasmic domain. The native signal peptide was replaced with a minimal version of the tissue plasminogen activator (mini-tPA) signal sequence (MDAMKRGLCCVLLLCGAVFVSPSAA). The extracellular HA domain was fused with a T4-foldon domain followed by a hexahistidine ($His_6$) tag to support formation of native-like trimers[1] and to facilitate purification, respectively. The coding sequence was RNA- and codon-optimized (Geneart / Thermo Fisher Scientific) and cloned into a derivative of pcDNA™5/FRT/TO vector. Expi293F cells were transiently transfected using the ExpiFectamine™ 293 Transfection Kit (Thermo Fisher Scientific), following the manufacturer's instructions.
Supernatant was cleared by centrifugation and loaded onto 5 mL HisTrap™ Excel column (Cytiva). Protein was eluted on an ÄKTA-Purifier system using a linear gradient elution to 0.5 M Imidazole. Peak fractions were pooled according to UV280 nm signal. Afterwards, the buffer was exchanged to PBS using desalting columns. To ensure homogeneity of native-like HA trimers, proteins were further purified using a Superdex 200 Increase 10/300 GL column. Peak fractions were collected and analyzed via blue native PAGE. Pooled trimer fractions were assessed for purity, oligomerization state and antigenicity using Blue Native PAGE (BN-PAGE), sodium dodecyl sulfate-PAGE (SDS-PAGE), size exclusion (SEC) and enzyme linked immunosorbent assay (ELISA) using HA-specific monoclonal antibodies CR9114[2] and 2D1[3].

#### Generation of soluble recombinant human CD40L for aptamer selection

Recombinant human CD40L (hCD40L; residues: 113–261 + C-terminal His-tag) used for aptamer selection was expressed and purified from BL21(DE3) *Escherichia coli* as described in Pedersen et al.[4]

#### Aptamer selection targeting Influenza hemagglutinin (HA) and human CD40 ligand (hCD40L):

For the selection of 2'F-CTP and LNA-TTP modified RNA aptamers targeting the Influenza A virus surface protein hemagglutinin (HA) and human CD40 ligand (hCD40L), the *N40-mU* RNA SELEX library was used. The library contained a 40-nt random region flanked by constant regions lacking uridines. Recombinant His-tagged, T4-foldon stabilized Influenza A H1N1 HA (A/England/195/09) and His-tagged hCD40L were immobilized on Dynabeads™ His-Tag

Isolation and Pulldown beads (Invitrogen, cat. no. 10103D) according to the manufacturer's protocol. For HA selection, 10, 7, 5, and 1 µg of protein were immobilized for rounds 1 to 4, respectively, while for hCD40L, the protein amounts were decreased as follows: 5 µg (round 1), 2 µg (round 2), 1 µg (rounds 3 and 4), 0.5 µg (rounds 5-7), 0.25 µg (rounds 8-9), and 0.125 µg (round 10).

Initial RNA libraries were generated via IVT as described above and refolded in 1x Dulbecco's Phosphate-Buffered Saline (D-PBS, Fisher Scientific, cat. no. 15217168) with 1 mM $MgCl_2$ using the following temperature program: 95 °C for 2 min, 65 °C for 5 min, 37 °C for 5 min, and held at 21 °C. The starting RNA amount for HA selection was 2000 pmol in round 1, decreasing to 200 pmol for rounds 2 to 4. For hCD40L selection, 1700 pmol RNA was used in round 1, reduced to 200 pmol in rounds 2 and 3, and kept at 150 pmol for rounds 4 to 10.

Selections were performed in SELEX buffer (1x D-PBS, 1 mM $MgCl_2$, 0.1 mg/mL BSA, 0.1 mg/mL salmon sperm DNA and for HA selection with 0.001% Tween20) at room temperature in a final volume of 300 µL for round 1 and 200 µL for rounds 2 to 4 for HA, while a constant volume of 200 µL was used for hCD40L. In rounds 2 to 4 for HA selection, the RNA libraries were precleared for 25 min with 3.5 µg His-tagged nanobody immobilized on Dynabeads™ to remove RNA with affinity for the His-tag and before target selection, they were also precleared for 20 min with empty beads. Incubation times were 45 min for rounds 1 and 2 and 30 min for rounds 3 and 4, with shaking at 750 rpm. For hCD40L selection, libraries were precleared with His-tagged nanobody beads for 25 min in all rounds except rounds 9 and 10, where two consecutive counter-selections were performed for 15 min each. The supernatant was then incubated with the target beads for 45 min for the initial rounds, decreasing to 10 min from round 4 onwards. After incubation, three washing steps were performed using 0.5 mL SELEX buffer for HA or 1x D-PBS with 1 mM $MgCl_2$ for hCD40L, with washing stringency increased over successive rounds.

Bound RNA was reverse transcribed, PCR amplified and purified as described above. The resulting DNA was used as a template for RNA transcription in the next selection round. RNA products were purified using the RNA Clean & Concentrator-25 kit.

**Preparation of sequencing libraries:** Selected DNA libraries for sequencing were first extended in a 50 µL PCR reaction using "Overhang fw" and "Overhang rev" primers. In a subsequent 30 µL PCR reaction, each library was further extended with unique indexes and sequencing adapters (Illumina Nextera XT Index Kit v2 Set A, cat. no. FC-131-2001). Both PCR products were purified using the GeneJET PCR Purification Kit. The sequencing libraries were then pooled in equal amounts, and free sequencing primers were removed using the Pippin Prep instrument with a 3% agarose cassette (Saga Science). Next-generation sequencing was performed on the iSeq 100 system (Illumina) using the iSeq 100 i1 Reagent v2 (300-cycle) kit (Illumina, cat. no. 20031371) following the manufacturer's protocol.

**NGS data analysis:** The preprocessing of the NGS data was consistent across all samples, while in-depth analysis varied depending on the sequenced sample.

***NGS data preprocessing:*** Reads were demultiplexed using the internal Illumina iSeq 100 software. Subsequent preprocessing was performed on the Galaxy server (https://usegalaxy.eu/) using single-read R1 reads. Quality control of FASTQ files was done using FastQC (v0.11.9; http://www.bioinformatics.babraham.ac.uk/projects/fastqc/). Adapter and constant regions were removed using Cutadapt (v4.0)[5] with default settings. Low-quality reads were removed using fastp (v0.23.2)[6] with qualified quality Phred set to 30. Reads collapsed to combine identical sequences and determine read counts. Data tables were exported and analyzed using custom Python (v3.10.9) scripts. A summary of the different analysis procedures is provided below, and the Python scripts can be obtained on request.

***Analysis of library diversity:*** Unique reads were quantified by dividing the total number of reads by the number of unique sequences. Sequences were filtered for a length of 40 nt, and only those sequences were considered for further analysis. Nucleotide distribution at each position and across the dataset was calculated, taking the number of reads into account. Base repeat frequencies were determined by grouping sequences by repeated bases and then aggregating the results across all sequences, factoring in the number of reads. Figures were generated using matplotlib (v3.7.0). Additional packages used were numpy (v1.23.5) and pandas (v1.5.3).

***Analysis of modified nucleotide incorporation fidelity:*** Sequences were filtered for a length of 40 nt, and only those sequences were considered for further analysis. Nucleotide distribution across all reads was calculated. The Levenshtein distance (LD) was used to match reads to one of the twelve reference sequences, with a threshold of LD = 10, to avoid multiple assignments. Substitutions were identified by comparing each read to its reference sequence, noting the sequence position, environment, and read count for each substitution detected. Substitutions were then grouped by patterns, base environments, and positions. Error rates were calculated per thousand bases by dividing the number of substitutions by the total number of nucleotides analyzed, considering the number of reads. Figures were generated using matplotlib (v3.7.0) and seaborn (Version 0.12.2). Further packages used were numpy (v1.23.5), pandas (v1.5.3), and for the LD, jellyfish (v0.9.0).

***Analysis of SELEX enrichment:*** For each SELEX experiment, preprocessed NGS data were combined, and similar sequences were clustered. Sequences were sorted by read number and filtered to include only those with more than 10 reads. The LD between sequences was calculated, starting with those having the highest read counts. Sequences with an LD < 5 were combined by summing their read counts and retaining the sequence with the higher read count. Enrichment for each sequence was calculated as a percentage of a respective pool by dividing its read count by the total read count of the pool. Exported tables displayed clustered sequences with their enrichment across analyzed SELEX rounds. Packages used were pandas (v1.5.3), and for the LD, jellyfish (v0.9.0).

**Transcription and purification of aptamers:** Selected clones were generated by IVT for characterization. The aptamers were transcribed with different pyrimidine bases in combination, following the transcription protocol described above. CTP, 2’F-CTP, UTP, dTTP were used at a final concentration of 2.5 mM, while LNA-dTTP was used at 1.25 mM. RNA was then purified either by denaturing PAGE or using the RNA Clean & Concentrator-25 kit as described above.

**HA aptamer biotinylation:** For affinity measurements, purified HA aptamers were biotinylated at the 3’-end by T4 ligation. For each RNA sample, a 50 µL reaction containing 1.5 mM ATP, 6 µM pCp-Biotin (Jena Bioscience, cat. no. NU-1706-BIO), 0.001 U/µL IPP, 0.3 U/µL T4 RNA ligase (Thermo Scientific, cat. no. EL0021), 15% PEG8000, 100 pmol RNA, and reaction buffer (50 mM Tris-HCl pH 7.5, 10 mM $MgCl_2$, 10 mM DTT) were incubated overnight at 4 °C. The biotinylated RNA was then purified using the RNA Clean & Concentrator-25 kit, and the total RNA concentration determined by UV-Vis spectroscopy.

**Affinity screening using biolayer interferometry:** The binding affinity of the selected aptamers and their variants with exchanged pyrimidine nucleotides toward their target proteins was measured using biolayer interferometry (BLI). All kinetic binding experiments were conducted in 96-well black flat- bottom plates (Greiner) using the Octet®Red96 system (ForteBio). Orbital shaking was set to 1,000 rpm for all BLI assays.

For aptamers targeting HA, biotinylated, folded RNA was immobilized on Octet® Streptavidin (SA) Biosensors (Sartorius, cat. no. 18-5019). The biosensors were then dipped into a series of 1/3 dilutions of T4-foldon stabilized HA trimer, starting from 300 nM, in BLI buffer (1x D-PBS, 1 mM MgCl2, 0.1 mg/mL BSA, 0.01% Tween20). The following experimental protocol was applied: baseline was generated in the BLI-containing well, followed by an association step (400 sec) in the protein solution wells, and a dissociation step (1200 sec) in the BLI buffer. A separate sensor was used for each HA concentration. For the screening of potential aptamers and determining the binding of aptamer variants, immobilized aptamers were tested with a constant concentration of 500 nM trimeric HA using the same protocol.

For aptamers targeting hCD40L, 2.5 µg/mL of His-tagged hCD40L protein diluted in BLI buffer was immobilized onto Octet® Ni-NTA Biosensors (Sartorius, cat. no. 18-5101). Serial dilutions of the aptamers (previously folded as described above) were prepared in binding buffer for the measurement. Baseline signals were recorded to establish initial BLI signals prior to each binding event (which included association, dissociation and regeneration steps. The protein-coated sensor was dipped into the aptamer solution for 400 sec (association step), followed by dipping into buffer alone for 1600 sec (dissociation step). Regeneration of the sensor was achieved by performing three cycles, each involving a 5 sec dip into glycine solution (10 mM at pH 1.4) followed by a 5 sec dip into buffer. For screening and determining the binding of the different aptamer variants, aptamers were tested at a constant concentration of 500 nM, following the same protocol described. The BLI data was analyzed using the instrument's software and Evilfit[7-8], with sensorgrams aligned to the baseline.

**Serum stability assay:** Gel-purified and refolded LNA aptamers and their variants were incubated at a concentration of 10 µM in 25% human serum in Dulbecco's Phosphate Buffered Saline (DPBS) at 37 °C. Samples were collected at various time points (0, 30 min, and 1, 3, 5, 24, 48, 72 and 96h). The collected samples were diluted 1:10 in MilliQ water, and 2 µL (equivalent to 2 pmol) were subjected to gel electrophoresis on an 8% denaturing polyacrylamide gel. The gels were quantified using ImageJ software (https://imagej.nih.gov/ij/).

**Melting temperatures of LNA aptamers:** UV measurements were performed on a Cary100 UV-Vis Spectrophotometer (Agilent Technologies) equipped with a temperature controller. A 500 µL solution of gel-purified and refolded LNA aptamers or their UTP version, each at a concentration of 1 µM, was prepared in 1x D-PBS containing 1 mM $MgCl_2$ and refolded as described previously. Buffer alone was used as a reference sample. Prior to the first melting curve scan, samples were held at 10 °C for 2 minutes. The temperature was then gradually increased to 90 °C at a rate of 0.2 °C per step. Subsequently, the temperature was lowered back to 10 °C following the same rate. In total, three heating curves from 10 °C to 90 °C and two cooling curves from 90 °C to 10 °C were recorded. The average of the two normalized (0 to 1) melting curves (from 10 °C to 90 °C) is shown. The first derivative of the averaged and normalized UV absorbance at 260 nm as a function of temperature was determined using OriginLab.

### Figure Section

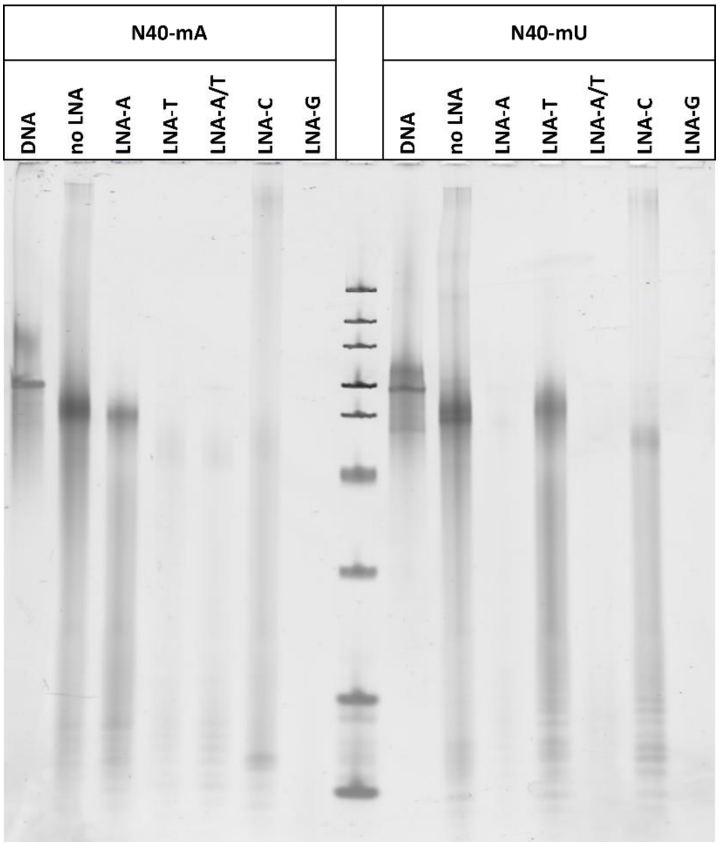

**Figure S1.** Urea-PAGE analysis of LNA-modified RNA. Transcribed RNA libraries (N40-mA and N40-mU) incorporating 2’F-pyrimidines and LNA-NTPs replacing corresponding nucleotides, alongside their respective DNA templates, were resolved on a 10% denaturing poly-acrylamide gel and visualized using SybrGold staining. An ultra-low range DNA ladder was used (band sizes from top: 300, 200, 150, 100, 75, 50, 35, 25, and 15 bp).

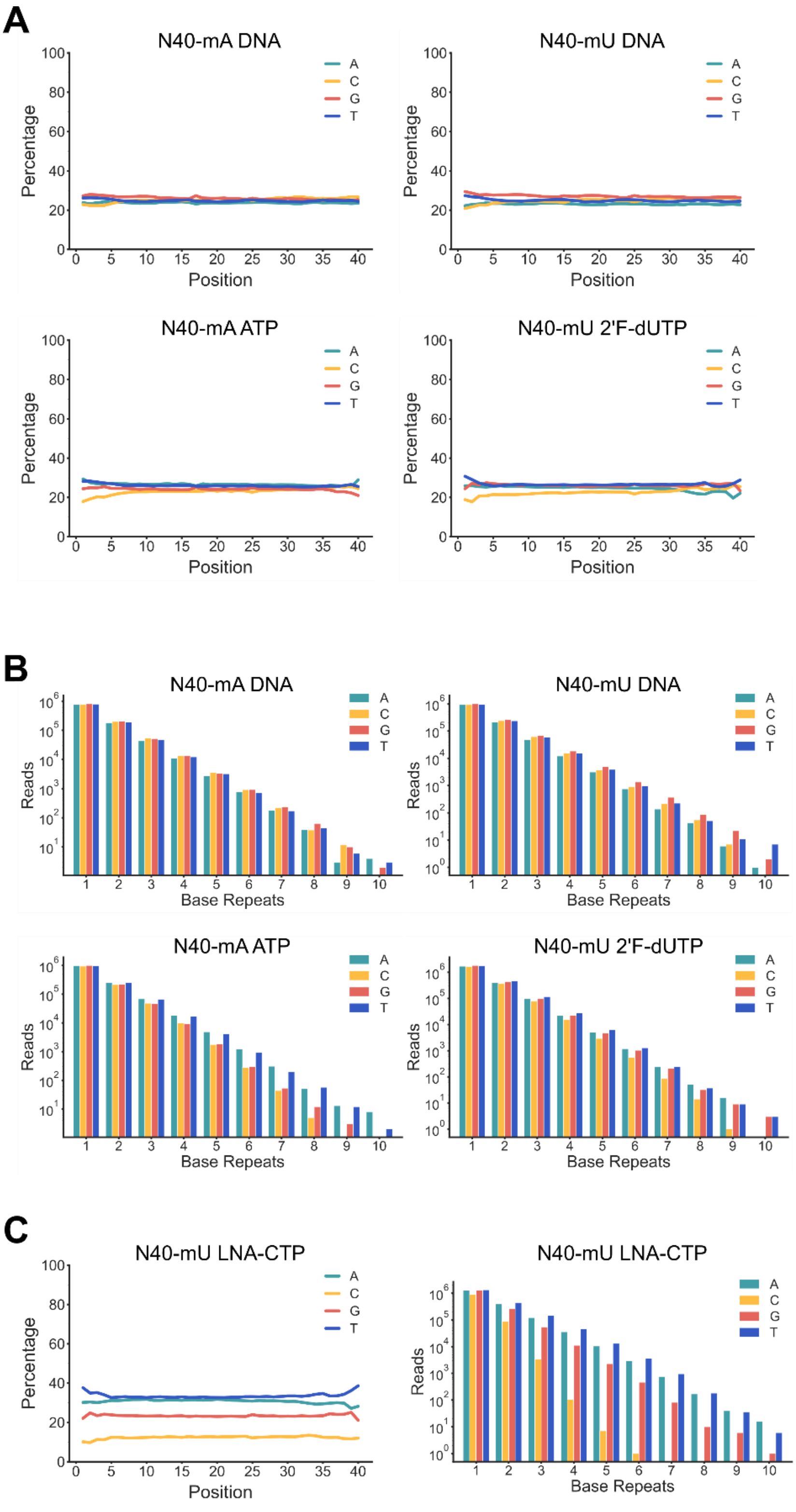

**Figure S2.** NGS analysis of LNA-modified RNA libraries. (A) Nucleotide distribution across the random region of the N40-mA and N40-mU DNA templates in reverse-transcribed control RNA libraries (N40-mA ATP, N40-mU 2’F-dUTP). (B) Abundance of reads containing repeats of the indicated number of bases in the sequenced libraries. (C) Nucleotide distribution and abundance of reads containing repeats of indicated number of bases of reverse transcribed LNA-CTP modified N40-mU RNA library.

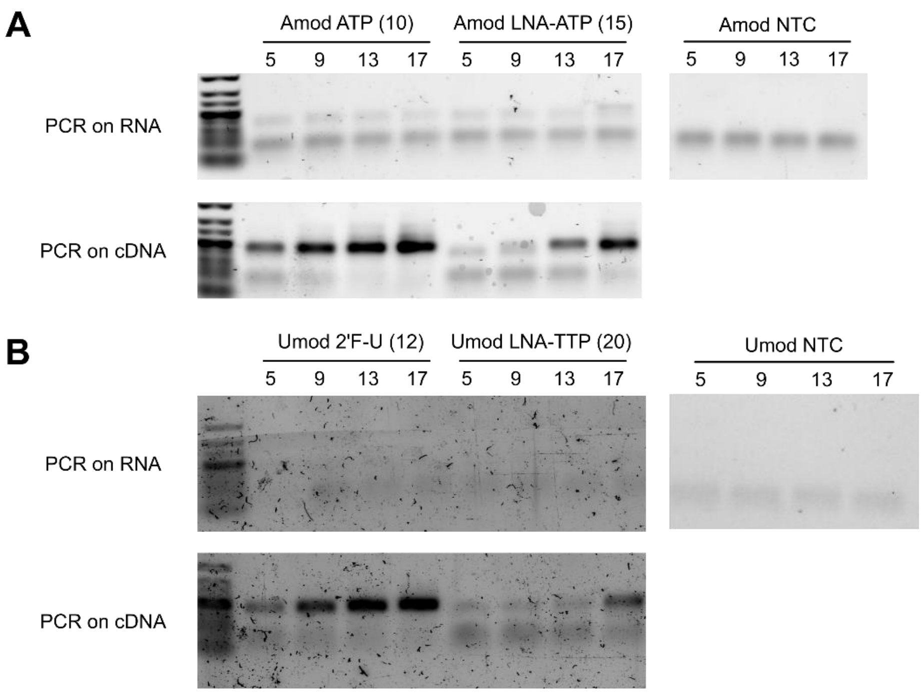

**Figure S3.** PCR analysis of RNA and cDNA. The RNA and cDNA of the (A) N40-mA and (B) N40-mU 12 sequences were PCR amplified to assess potential residual DNA template after DNase I treatment and RNA purification. PCR samples were collected after 5, 9, 13, and 17 amplification cycles and visualized in a 3% agarose gel stained with SybrSafe. The number in brackets indicates amplification cycles used for the final PCR of the sample processed for NGS. NTC = non-template control. An ultra-low range DNA ladder was used (Band sizes from top: 300, 200, 150, 100, 75, 50, 35, 25, and 15 bp).

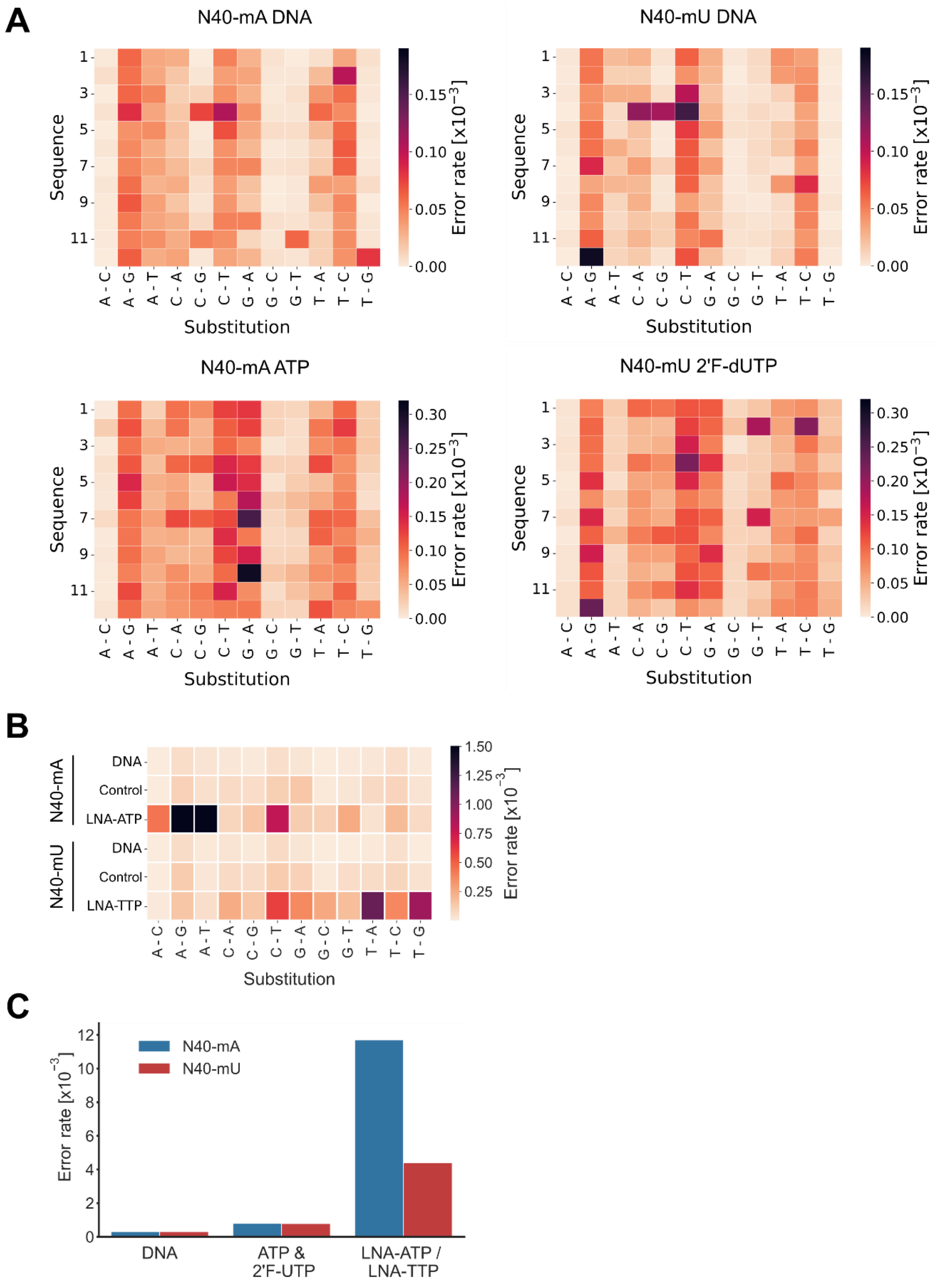

**Figure S4.** NGS analysis of LNA incorporation into the 12 test sequences. (A) Substitution error rates for N40-mA and N40-mU DNA templates and reverse transcribed control RNAs (N40-mA ATP and N40-mU 2’F-dUTP) across the 12 tested sequences. Heatmap displaying combined error rates for each sequence and observed substitution patterns (expected base - detected base). (B) Heatmap summarizing combined error rates for each substitution pattern (expected base - detected base) across all 12 sequences for DNA templates, control IVT, and LNA-modified IVT samples. For N40-mA (LNA-ATP), the A-G (6.7 *$10^{-3}$) and A-T (2.7 *$10^{-3}$) substitution error rates exceed the displayed scale. (C) Comparison of combined error rates for the DNA templates, control RNAs (N40-mA and N40-mU transcribed with 2’F-modified pyrimidines), and LNA-modified RNA libraries. All error rates are relative to the number of sequenced nucleotides.

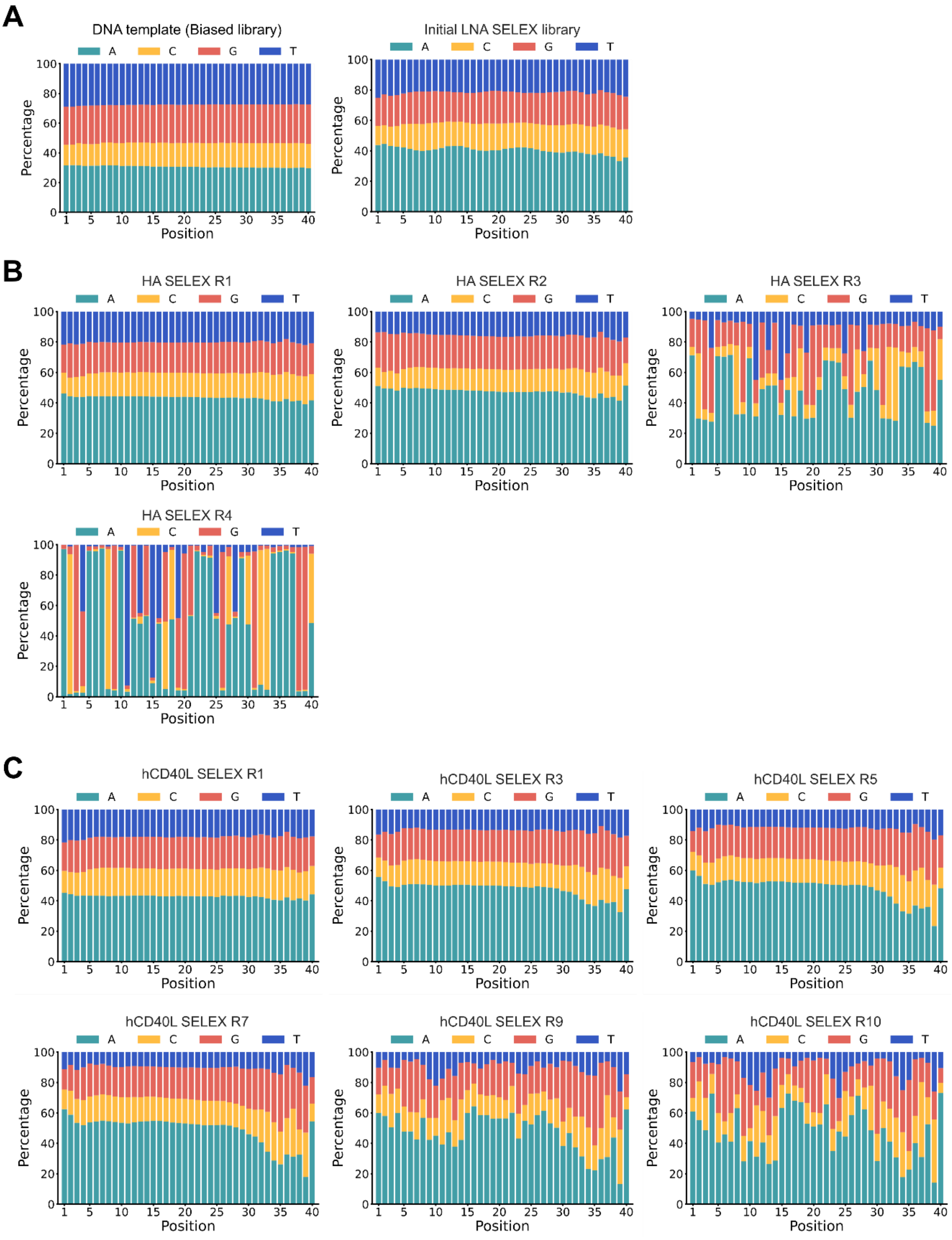

**Figure S5.** Nucleotide distribution across the random region in SELEX pools. Nucleotide distribution in the sequenced DNA pools from the (A) N40-mU SELEX library DNA template and initial LNA-modified RNA SELEX library, and of selected pools of the (B) HA-targeted and the (C) hCD40L-targeted SELEX. The initial LNA-SELEX library was generated from a DNA template library synthesized by Integrated DNA Technologies (IDT), which contained a lower overall cytosines content and a higher adenines frequency relative to the other bases.

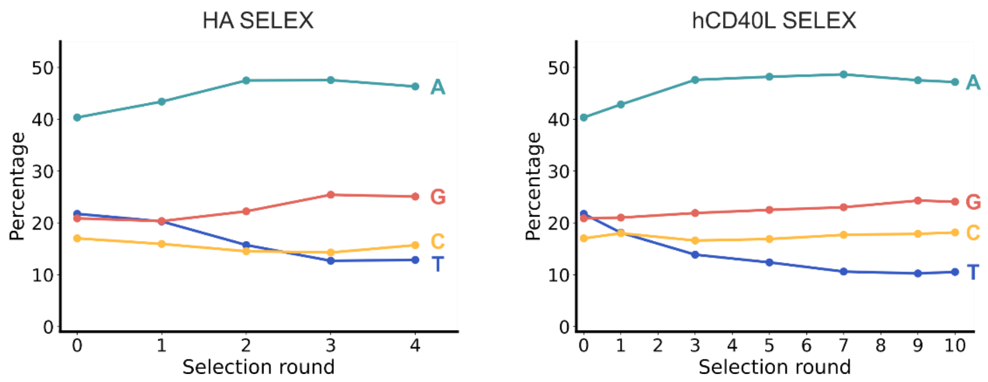

**Figure S6.** Overall nucleotide distribution in sequenced DNA pools from the different SELEX rounds.

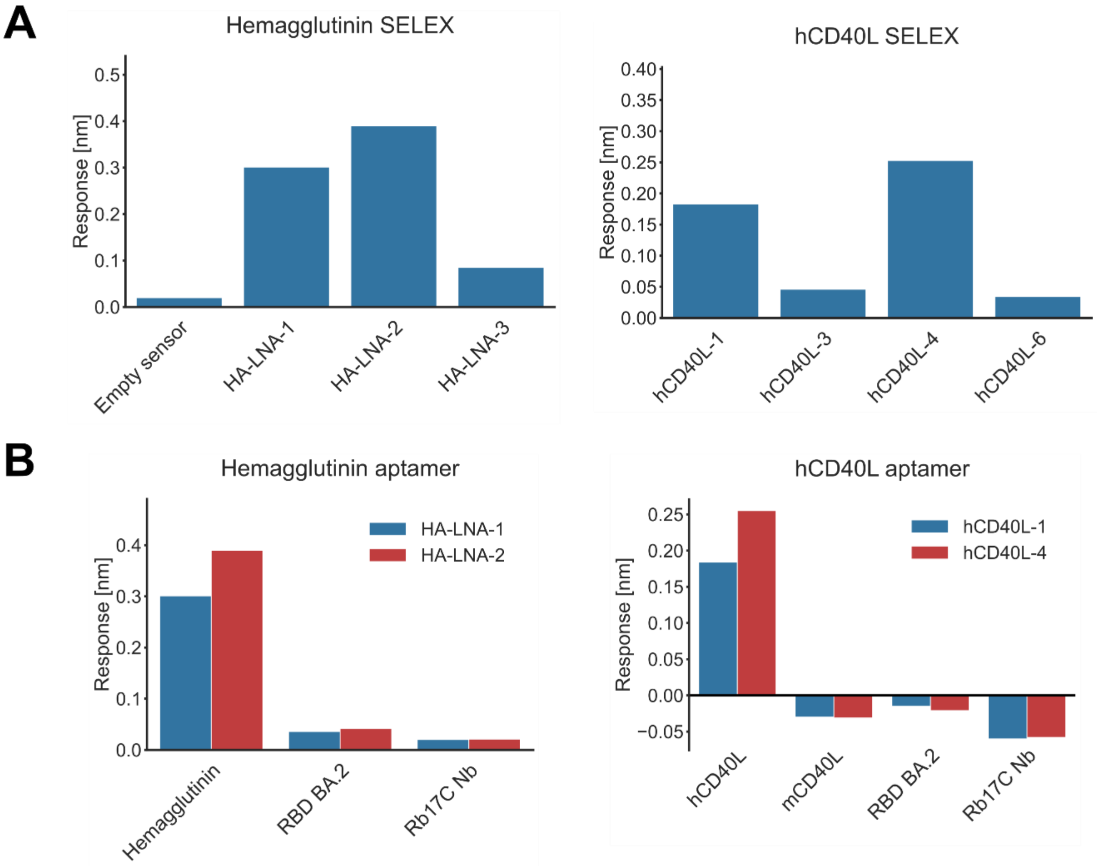

**Figure S7.** BLI screening for aptamer binding and specificity. (A) Screening of selected RNA sequences from the HA- and hCD40L-targeted selections for target binding. (Left) Biotinylated RNA was immobilized on streptavidin-coated sensors and exposed to 500 nM HA trimer. (Right) His-tagged hCD40L was immobilized on Ni-NTA-coated sensors and exposed to 500 nM RNA (B) Specificity of selected aptamers for their target (HA or hCD40L) was assessed against unrelated His-tagged proteins (SarS-CoV-2 receptor binding domain, RBD and Rb17C nanobody) under the same conditions as in A. For A and B, the response after 400 sec of association is shown.

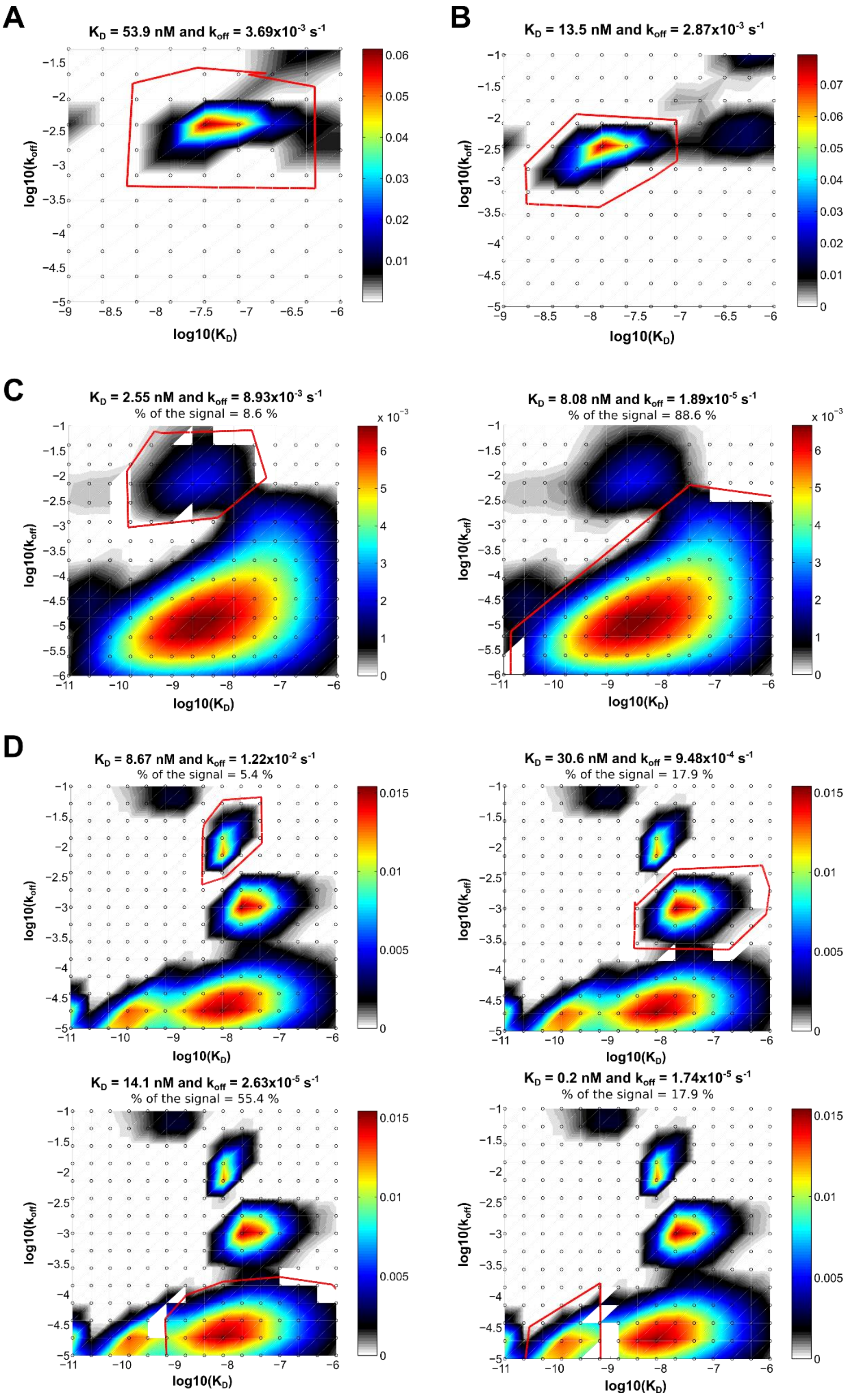

**Figure S8.** Modeled distributions of binding kinetics. For (A) HA-LNA-1, (B) HA-LNA-2, (C) hCD40L-1, and (D) hCD40L-4), the modeled distribution of binding kinetics from the BLI data is shown, obtained using Evilfit[7-8] analysis. The 2D plots display the distributions as contour maps with $log_{10}$(KD) and $log_{10}$(koff) on the x- and y-axes, respectively. The scale bar represents signal intensity.

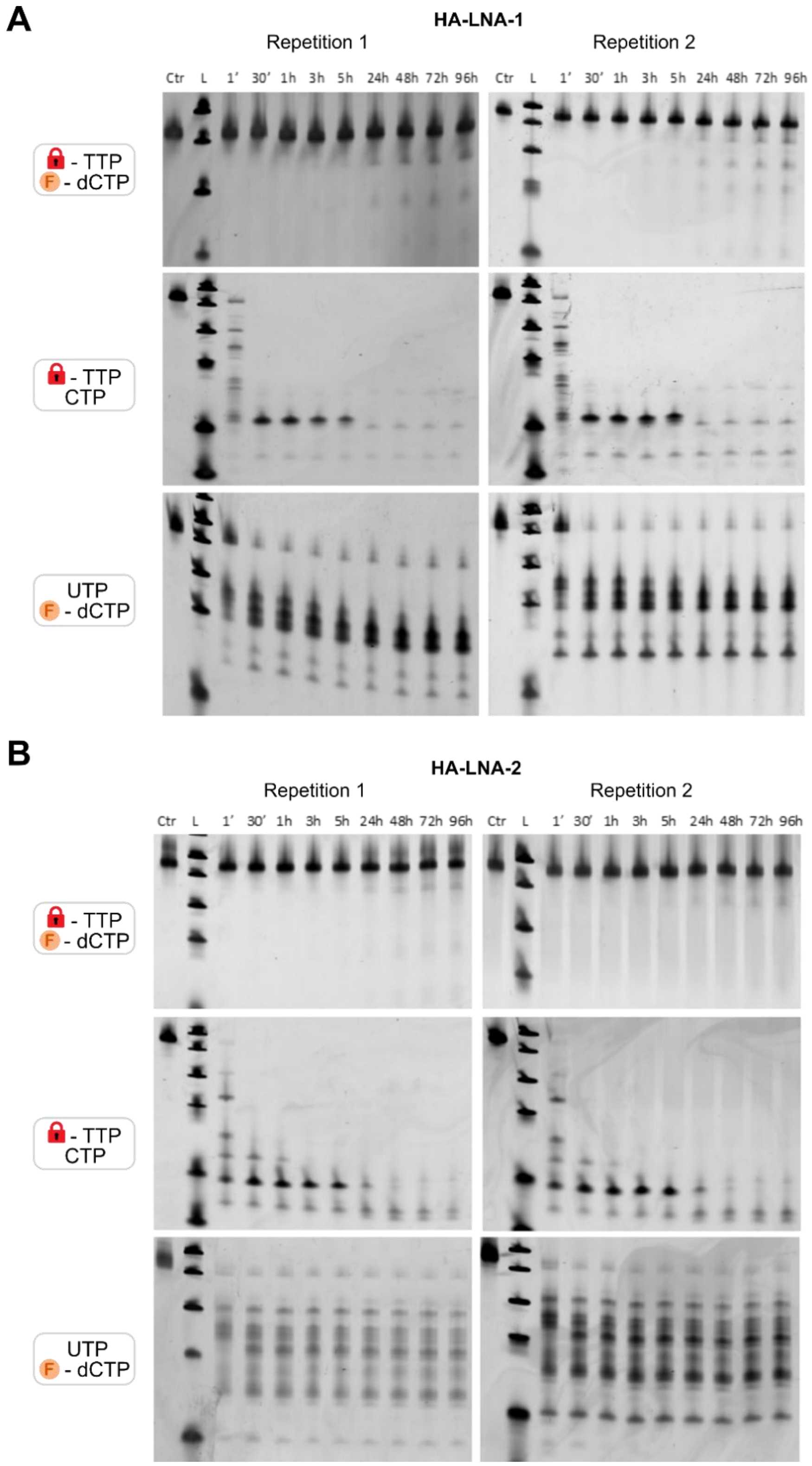

**Figure S9.** Aptamer serum stability analyzed on denaturing PAGE. RNA of (A) HA-LNA-1 and (B) HA-LNA-2 with different chemical modifications after incubation in 25% human plasma over four days, visualized on denaturing PAGE and stained with SybrGold. Two replicates are shown for each sample. The control sample was not incubated with serum. The second lane contains the ultra-low range DNA ladder (Band sizes from top: 100, 75, 50, 35, 25, and 15 bp).

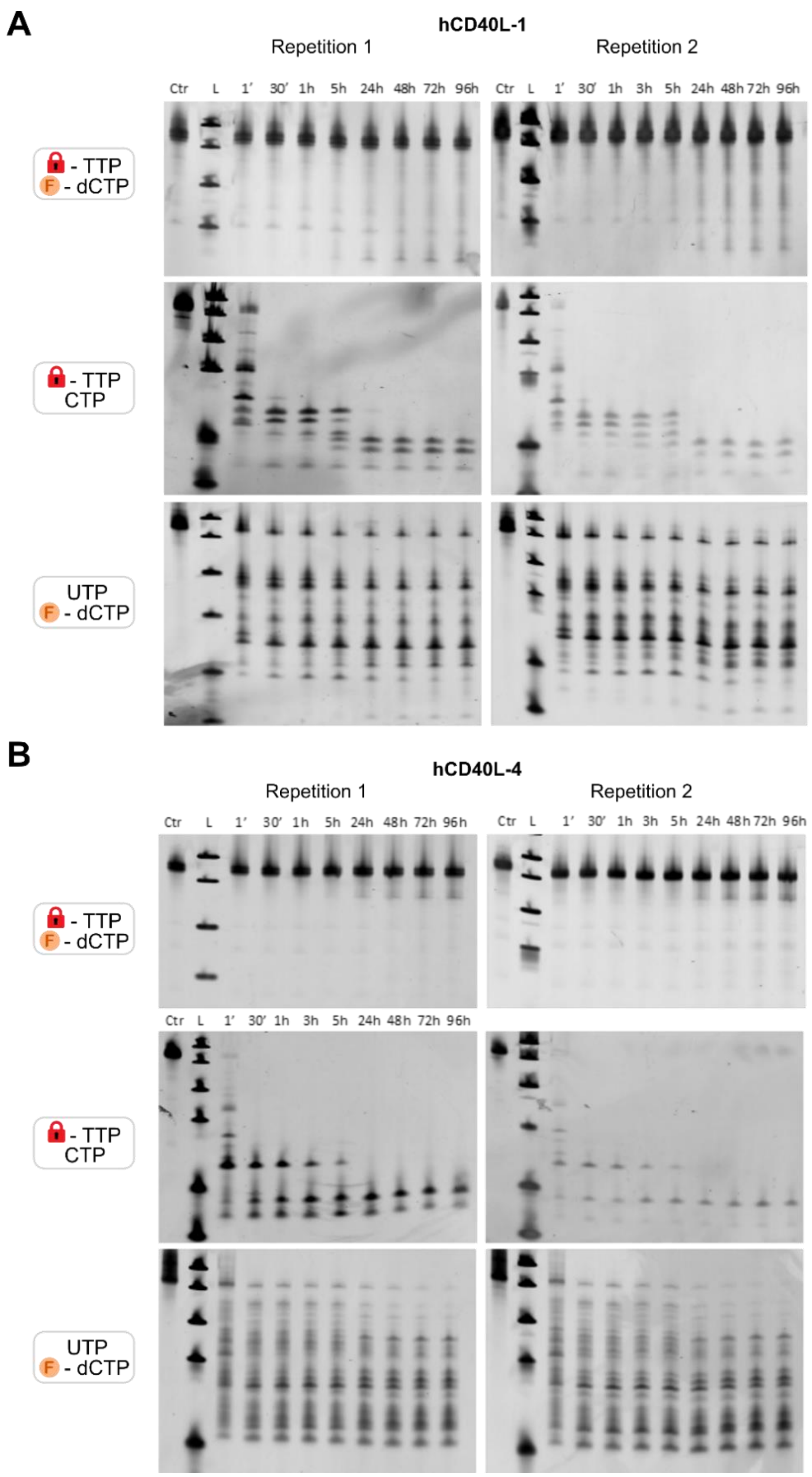

**Figure S10.** Aptamer serum stability analyzed on denaturing PAGE. RNA of (A) hCD40L-1 and (B) hCD40L-4 with different chemical modifications after incubation in 25% human plasma over four days, visualized on denaturing PAGE and stained with SybrGold. Two replicates are shown for each sample. The control sample was not incubated with serum. The second lane contains the ultra-low range DNA ladder (Band sizes from top: 100, 75, 50, 35, 25, and 15 bp).

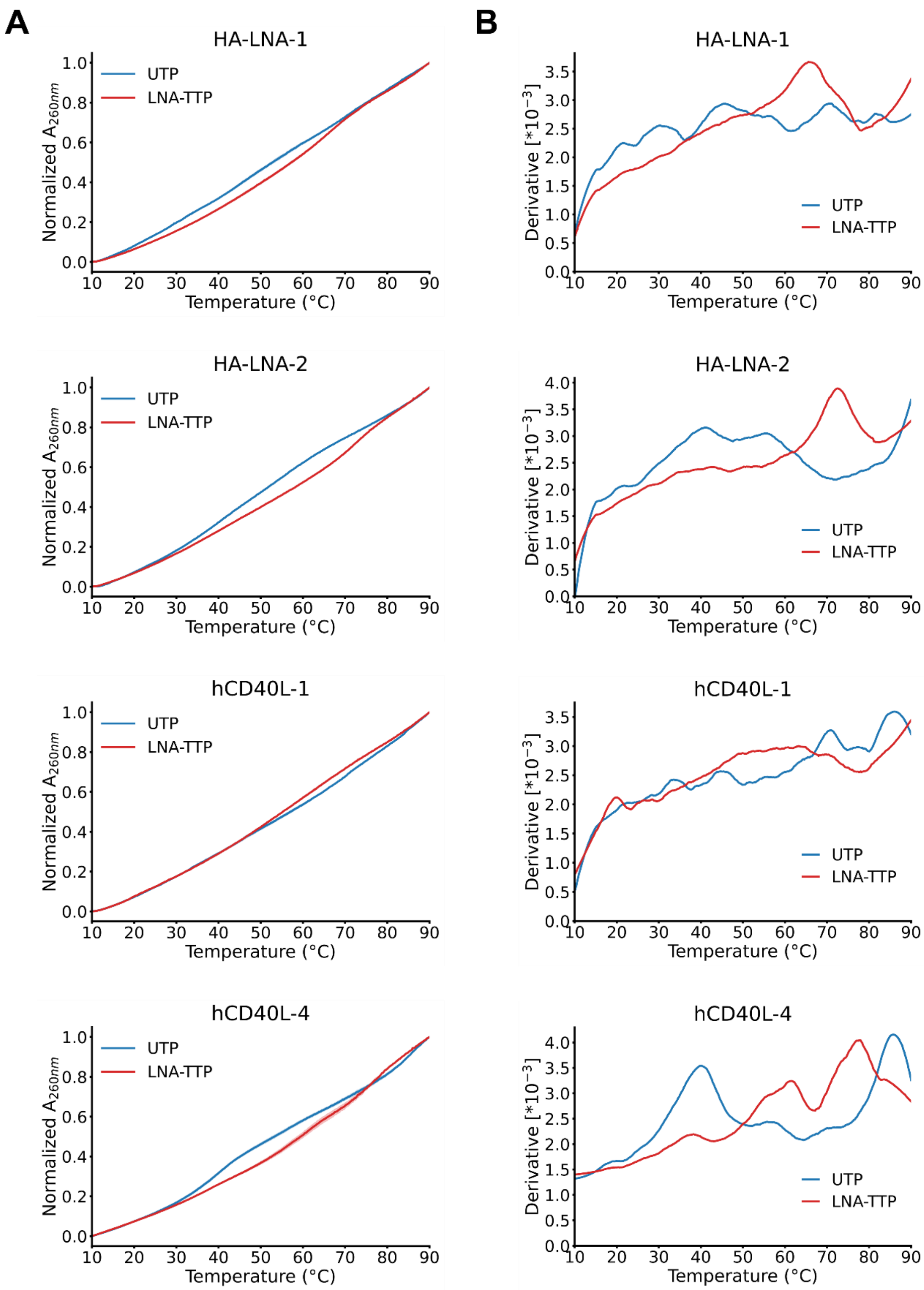

**Figure S11.** Thermal stability of LNA- and 2’F-modified RNA aptamers. (A) Normalized melting curves of RNA aptamers transcribed with 2'F-dCTP and either LNA-TTP or UTP, recorded by monitoring absorbance at 260 nm from 10 °C to 90 °C. Data represent the average of two measurements, with standard deviation indicated as shaded background. (B) First derivate of the averaged melting curves, highlighting the melting transitions in the melting curves.

### Table Section

**Table S1.** Design of the libraries and the twelve sequences used for NGS analysis of LNA incorporation.

| | |
|---|---|
| **Constant regions** | |
| **5'-N40-mA** | GGGUGGUCGUUGCGC |
| **3'-N40-mA** | CUGUCGUCUGGCGCGU |
| **5'-N40-mU** | GGGAGGAGGAAGCGG |
| **3'-N40-mU** | CAGACGACACGCCCGA |
| **N40 libraries RNA** | |
| **N40-mA** | GGGUGGUCGUUGCGC-N40-CUGUCGUCUGGCGCGU |
| **N40-mU** | GGGAGGAGGAAGCGG-N40-CAGACGACACGCCCGA |
| **Random regions of 12 N40-mA sequences** | |
| **A1** | ACGCUAUAUACAAUUCUUCGGUCAGGCCGUGAAUCGGCGA |
| **A2** | AGCUUGCUCGGACAACCAGUUGCCAUUGGAAAUGCAUUCG |
| **A3** | AGGGGAACUGUCCGGGAUUUCCAAUCCUGAACUACAUUCG |
| **A4** | CGCCCGCUAUUUAGGGUCUAAAAUGGGUGUCAUCAACCGA |
| **A5** | GGUCCGCGAUUUUACACCCGGCUUUGGAUAAGGCACAUAA |
| **A6** | GUAACGUGGGUACUCAACAAGCCACCUUUUAGGUGCCAUG |
| **A7** | GUCGACGUUCUCAGAAGCUGUGAACCGCCAUAGCUUAUAG |
| **A8** | UACGUCCUUGUGUGACGAAAGGAACUCCAUGGUGCAACCU |
| **A9** | UAGCUUCAUGAGCGACUGAUCCGGGGAACUUCGAUCUCAA |
| **A10** | UCACGCACUAACCUUUAUUGAUGCCAUGAACGGAGCUGGG |
| **A11** | UGGUCUUUGGAGGCGAAGACCACACAUGUUUUCGACCCAA |
| **A12** | UGUUCUUCAACUGGCUGCAGAACCGCAACUGAGCGUUGAA |
| **Random regions of 12 N40-mU sequences** | |
| **U1** | UCGCAUAUAUCUUAACAACGGACUGGCCGAGUUACGGCGU |
| **U2** | UGCAAGCACGGUCUUCCUGAAGCCUAAGGUUUAGCUAACG |
| **U3** | UGGGGUUCAGACCGGGUAAACCUUACCAGUUCAUCUAACG |
| **U4** | CGCCCGCAUAAAUGGGACAUUUUAGGGAGACUACUUCCGU |
| **U5** | GGACCGCGUAAAAUCUCCCGGCAAAGGUAUUGGCUCUAUU |
| **U6** | GAUUCGAGGGAUCACUUCUUGCCUCCAAAAUGGAGCCUAG |
| **U7** | GACGUCGAACACUGUUGCAGAGUUCCGCCUAUGCAAUAUG |
| **U8** | AUCGACCAAGAGAGUCGUUUGGUUCACCUAGGAGCUUCCA |
| **U9** | AUGCAACUAGUGCGUCAGUACCGGGGUUCAACGUACACUU |
| **U10** | ACUCGCUCAUUCCAAAUAAGUAGCCUAGUUCGGUGCAGGG |
| **U11** | AGGACAAAGGUGGCGUUGUCCUCUCUAGAAAACGUCCCUU |
| **U12** | AGAACAACUUCAGGCAGCUGUUCCGCUUCAGUGCGAAGUU |

**Table S2.** Statistics for NGS analysis of LNA incorporation into N40-mA and N40-mU libraries.

| Modification | Library | Total reads | Unique Reads | Nucleotide [%] (40 nt length) | | | | Length [%] | | |
|---|---|---|---|---|---|---|---|---|---|---|
| | | | | A | C | G | T | < 40 nt | 40 nt | > 40 nt |
| ATP | N40-mA | 185682 | 174900 | 26.7 | 23.1 | 24.1 | 26.1 | 4.8 | 91.7 | 3.6 |
| LNA-ATP | N40-mA | 190177 | 173805 | 18.3 | 20.4 | 22.2 | 39.2 | 14.3 | 80.9 | 4.9 |
| 2'F-UTP | N40-mU | 321390 | 292478 | 24.5 | 22.6 | 26.2 | 26.7 | 2.7 | 93.3 | 4.0 |
| LNA-TTP | N40-mU | 214798 | 196007 | 34.6 | 22.1 | 23.0 | 20.3 | 9.2 | 85.7 | 5.1 |
| LNA-CTP | N40-mU | 239403 | 196080 | 30.7 | 12.3 | 23.4 | 33.6 | 5.6 | 90.9 | 3.4 |
| DNA | N40-mA | 149742 | 125820 | 23.7 | 25.3 | 26.1 | 24.9 | 3.6 | 93.2 | 3.3 |
| DNA | N40-mU | 180086 | 155984 | 23.1 | 24.8 | 27.1 | 25.0 | 2.5 | 94.3 | 3.3 |

**Table S3.** Detailed statistics for NGS analysis of LNA incorporation into twelve test sequences from the N40-mA and N40-mU libraries, including error analysis for each individual sequence.

| N40-mA | | | | | | | | | |
|---|---|---|---|---|---|---|---|---|---|
| | DNA | | | Control | | | LNA-ATP (N40-mA) | | |
| Clone | Reads | Reads [%] | Error rate [$10^{-3}$] | Reads | Reads [%] | Error rate [$10^{-3}$] | Reads | Reads [%] | Error rate [$10^{-3}$] |
| **1** | 33436 | 8.98 | 0.25 | 22912 | 7.33 | 0.77 | 16198.00 | 6.69 | 12.90 |
| **2** | 35089 | 9.43 | 0.33 | 26043 | 8.33 | 0.84 | 16496.00 | 6.82 | 20.12 |
| **3** | 37974 | 10.20 | 0.32 | 26973 | 8.63 | 0.63 | 17613.00 | 7.28 | 18.21 |
| **4** | 22068 | 5.93 | 0.48 | 10384 | 3.32 | 0.91 | 27392.00 | 11.32 | 9.88 |
| **5** | 27142 | 7.29 | 0.31 | 19818 | 6.34 | 0.75 | 6413.00 | 2.65 | 17.78 |
| **6** | 32302 | 8.68 | 0.25 | 47625 | 15.24 | 0.75 | 14310.00 | 5.91 | 11.44 |
| **7** | 27015 | 7.26 | 0.31 | 13580 | 4.35 | 1.07 | 17472.00 | 7.22 | 9.07 |
| **8** | 32552 | 8.74 | 0.27 | 35694 | 11.42 | 0.73 | 5543.00 | 2.29 | 12.96 |
| **9** | 30067 | 8.08 | 0.26 | 19903 | 6.37 | 0.75 | 30330.00 | 12.53 | 7.36 |
| **10** | 32290 | 8.67 | 0.29 | 33977 | 10.87 | 0.93 | 28072.00 | 11.60 | 10.02 |
| **11** | 30722 | 8.25 | 0.36 | 27729 | 8.87 | 0.80 | 18413.00 | 7.61 | 13.48 |
| **12** | 31595 | 8.49 | 0.34 | 27889 | 8.92 | 0.80 | 43755.00 | 18.08 | 10.01 |

| N40-mU | | | | | | | | | |
|---|---|---|---|---|---|---|---|---|---|
| | DNA | | | Control | | | LNA-TTP (N40-mU) | | |
| Clone | Reads | Reads [%] | Error rate [$10^{-3}$] | Reads | Reads [%] | Error rate [$10^{-3}$] | Reads | Reads [%] | Error rate [$10^{-3}$] |
| **1** | 27633 | 9.46 | 0.30 | 33367 | 10.31 | 0.77 | 31960 | 16.98 | 5.34 |
| **2** | 22371 | 7.66 | 0.24 | 25082 | 7.75 | 0.96 | 5746 | 3.05 | 3.85 |
| **3** | 20639 | 7.06 | 0.31 | 18793 | 5.81 | 0.67 | 33830 | 17.97 | 2.46 |
| **4** | 26899 | 9.21 | 0.53 | 31481 | 9.72 | 0.83 | 119 | 0.06 | 16.81 |
| **5** | 25379 | 8.69 | 0.28 | 13567 | 4.19 | 0.77 | 13204 | 7.02 | 3.75 |
| **6** | 17632 | 6.03 | 0.27 | 17570 | 5.43 | 0.58 | 5739 | 3.05 | 4.01 |
| **7** | 19513 | 6.68 | 0.31 | 29595 | 9.14 | 0.88 | 27493 | 14.61 | 2.89 |
| **8** | 24310 | 8.32 | 0.30 | 21095 | 6.52 | 0.78 | 4842 | 2.57 | 4.37 |
| **9** | 28306 | 9.69 | 0.21 | 39362 | 12.16 | 0.76 | 8473 | 4.50 | 2.78 |
| **10** | 26423 | 9.04 | 0.22 | 30229 | 9.34 | 0.74 | 16373 | 8.70 | 4.69 |
| **11** | 25661 | 8.78 | 0.29 | 29768 | 9.20 | 0.78 | 17672 | 9.39 | 6.55 |
| **12** | 27436 | 9.39 | 0.38 | 33823 | 10.45 | 0.74 | 22756 | 12.09 | 7.08 |

**Table S4.** Statistics for NGS analysis of LNA incorporation into 12 test sequences from the N40-mA and N40-mU libraries. Unassigned sequences could not be assigned to any of the twelve test sequences.

| Sample | Library | Total reads | Nucleotide [%] (40 nt length) | | | | Length [%] | | | Reads with X Errors [%] | | | | Overall Error rate [$10^{-3}$] | Unassigned sequences [%] | | Insertion/ Deletion [%] |
|---|---|---|---|---|---|---|---|---|---|---|---|---|---|---|---|---|---|
| | | | A | C | G | T | < 40 nt | 40 nt | > 40 nt | 0 | 1 | 2 | 3 | | Reads | Sequence | |
| ATP | N40-mA | 323200 | 25.1 | 24.8 | 25.1 | 25.0 | 2.0 | 96.7 | 1.3 | 96.98 | 2.88 | 0.12 | 0.01 | 0.79 | 0.01 | 1.02 | 0.014 |
| LNA-ATP | N40-mA | 275626 | 21.1 | 24.9 | 27.2 | 26.8 | 10.8 | 87.9 | 1.3 | 64.80 | 26.04 | 7.12 | 1.70 | 11.70 | 0.05 | 0.54 | 0.047 |
| 2'F-UTP | N40-mU | 340607 | 25.2 | 24.7 | 25.1 | 25.0 | 3.1 | 95.1 | 1.8 | 97.07 | 2.78 | 0.14 | 0.01 | 0.78 | 0.00 | 0.13 | 0.026 |
| LNA-TTP | N40-mU | 254137 | 28.5 | 23.9 | 25.0 | 22.7 | 19.1 | 77.3 | 3.6 | 85.39 | 12.45 | 1.75 | 0.27 | 4.40 | 4.03 | 9.74 | 0.098 |
| DNA | N40-mA | 380621 | 24.9 | 24.8 | 25.2 | 25.2 | 1.6 | 97.8 | 0.6 | 98.84 | 1.10 | 0.05 | 0.01 | 0.31 | 0.00 | 0.36 | 0.007 |
| DNA | N40-mU | 302513 | 25.0 | 24.7 | 25.0 | 25.3 | 2.8 | 96.6 | 0.7 | 98.89 | 1.02 | 0.07 | 0.01 | 0.31 | 0.00 | 0.10 | 0.008 |

**Table S5.** Top 10 most abundant sequences in HA- and hCD40L-targeted SELEX experiments after four and ten selection rounds, respectively. The random region of the aptamer sequences is shown. (*) Sequences generated by IVT and tested in BLI screening. Common motifs identified using the MEME tool[9] are highlighted by different colors.

| Target | Name | Frequency [%] | Number of Ts | Sequence |
|---|---|---|---|---|
| HA | *HA-LNA-1 | 43.24 | 3 | ACGGAAACGATGAATTGCGGGAAAAGCAAAGCCAAAAGGC |
| | *HA-LNA-2 | 42.82 | 7 | ACGTAAACGATATGTACATGAAAATGATACGCCAAAAGGA |
| | *HA-LNA-3 | 3.95 | 9 | ACGCAAACGATACAATTATTAATGATGATTGACAAAAGGG |
| | HA-LNA-4 | 0.89 | 9 | GTAATCAAGATGTACAACATAGAACCAGGATGATAATTAC |
| | HA-LNA-5 | 0.84 | 6 | AGCGGTATATCGAAAGATCAAAACTATAACAAAGGAGAAA |
| | HA-LNA-6 | 0.17 | 1 | AGGGAAGAGAGAGGAAACGAAGAGAACGGGACGATGAGGA |
| | HA-LNA-7 | 0.06 | 5 | GGGGAGAAAAGGACGCTCGTAAATATCAAAAAGACGGATA |
| | HA-LNA-8 | 0.05 | 1 | AGGGAAGGAAAACAAAAACGGGGTGACACGCCACAGAAGA |
| | HA-LNA-9 | 0.04 | 3 | AGGGAAAAAGGACCTGTCGAAGAAGAAAAGAGAAATAAGC |
| | HA-LNA-10 | 0.04 | 8 | AAAAAGAATTGCAGAAGTTGAACTCGAAAGACTTCAAGTG |
| hCD40L | *hCD40L-1 | 11.58 | 4 | AAAAAGCAGGTGTGAAAAGGACGTCAAACGACGTGACACA |
| | hCD40L-2 | 6.49 | 5 | GCGAGAAAGGCATGAAAAGGGATTCAAACGAATCAACATA |
| | *hCD40L-3 | 6.37 | 2 | GGAACAGGGATTCCCAAAAAGAGAAGAAAGCGCGGGCACA |
| | *hCD40L-4 | 3.72 | 8 | ACGATGAACAAGTTGAAAAGGAATTTAAACGAATTAACAA |
| | hCD40L-5 | 2.80 | 4 | GCAAAAAAACAGCAGAAGTAACGATGCCAGAAAGATGGCAGTA |
| | *hCD40L-6 | 2.72 | 5 | AAAACGTACAAAATAGGGCAAAGTGAACAGGGATTCACGA |
| | hCD40L-7 | 2.42 | 4 | CAACAGGGATTGCCAACAAAAAAATACACGAAACGGACGT |
| | hCD40L-8 | 2.03 | 5 | AACAGGGATTCCGGAAAGAAAATAAATGACGGGCTGAACA |
| | hCD40L-9 | 1.22 | 3 | AACAGGGATTCCGAAATAAAACAAAAACAAAACGGGCAGG |
| | hCD40L-10 | 1.21 | 6 | AAATAAGTAACAGGGATTACAACAAAATAAGGGCACAGTG |

**Table S6.** Sequenced synthesized by Integrated DNA Technologies (IDT).

| Name | Sequence | Length |
|---|---|---|
| N40-mU library | TCGGGCGTGTCGTCTG-N40-CCGCTTCCTCCTCCC | 71 |
| N40-mA library | ACGCGCCAGACGACAG-N40-GCGCAACGACCACCC | 71 |
| N40-mU forward primer (5') | TAATACGACTCACTATAGGGAGGAGGAAGCGG | 32 |
| N40-mU reverse primer (3') | TCGGGCGTGTCGTCTG | 16 |

| | | |
|---|---|---|
| N40-mA forward primer (5') | TAATACGACTCACTATAGGGTGGTCGTTGCGC | 32 |
| JL40 Amod reverse primer (3') | ACGCGCCAGACGACAG | 16 |
| N40-mU NGS overhang forward primer | *TCGTCGGCAGCGTCAGATGTGTATAAGAGACAG*TAATACGACTCACTATAGGGAGGAGGAAGCGG | 65 |
| N40-mU NGS overhang reverse primer | *GTCTCGTGGGCTCGGAGATGTGTATAAGAGACAG*TCGGGCGTGTCGTCTG | 50 |
| N40-mA NGS overhang forward primer | *TCGTCGGCAGCGTCAGATGTGTATAAGAGACAG*TAATACGACTCACTATAGGGTGGTCGTTGCGC | 65 |
| N40-mA NGS overhang reverse primer | *GTCTCGTGGGCTCGGAGATGTGTATAAGAGACAG*ACGCGCCAGACGACAG | 50 |
| **<u>ssDNA for 12 clones</u>** | | |
| **N40-mU** | | |
| U1 | TCGGGCGTGTCGTCTGACGCCGTAACTCGGCCAGTCCGTTGTTAAGATATATGCGACCGCTTCCTCCTCCC | 71 |
| U2 | TCGGGCGTGTCGTCTGCGTTAGCTAAACCTTAGGCTTCAGGAAGACCGTGCTTGCACCGCTTCCTCCTCCC | 71 |
| U3 | TCGGGCGTGTCGTCTGCGTTAGATGAACTGGTAAGGTTTACCCGGTCTGAACCCCACCGCTTCCTCCTCCC | 71 |
| U4 | TCGGGCGTGTCGTCTGACGGAAGTAGTCTCCCTAAAATGTCCCATTTATGCGGGCGCCGCTTCCTCCTCCC | 71 |
| U5 | TCGGGCGTGTCGTCTGAATAGAGCCAATACCTTTGCCGGGAGATTTTACGCGGTCCCCGCTTCCTCCTCCC | 71 |
| U6 | TCGGGCGTGTCGTCTGCTAGGCTCCATTTTGGAGGCAAGAAGTGATCCCTCGAATCCCGCTTCCTCCTCCC | 71 |
| U7 | TCGGGCGTGTCGTCTGCATATTGCATAGGCGGAACTCTGCAACAGTGTTCGACGTCCCGCTTCCTCCTCCC | 71 |
| U8 | TCGGGCGTGTCGTCTGTGGAAGCTCCTAGGTGAACCAAACGACTCTCTTGGTCGATCCGCTTCCTCCTCCC | 71 |
| U9 | TCGGGCGTGTCGTCTGAAGTGTACGTTGAACCCCGGTACTGACGCACTAGTTGCATCCGCTTCCTCCTCCC | 71 |
| U10 | TCGGGCGTGTCGTCTGCCCTGCACCGAACTAGGCTACTTATTTGGAATGAGCGAGTCCGCTTCCTCCTCCC | 71 |
| U11 | TCGGGCGTGTCGTCTGAAGGGACGTTTTCTAGAGAGGACAACGCCACCTTTGTCCTCCGCTTCCTCCTCCC | 71 |
| U12 | TCGGGCGTGTCGTCTGAACTTCGCACTGAAGCGGAACAGCTGCCTGAAGTTGTTCTCCGCTTCCTCCTCCC | 71 |
| **N40-mA** | | |
| A1 | ACGCGCCAGACGACAGTCGCCGATTCACGGCCTGACCGAAGAATTGTATATAGCGTGCGCAACGACCACCC | 71 |
| A2 | ACGCGCCAGACGACAGCGAATGCATTTCCAATGGCAACTGGTTGTCCGAGCAAGCTGCGCAACGACCACCC | 71 |
| A3 | ACGCGCCAGACGACAGCGAATGTAGTTCAGGATTGGAAATCCCGGACAGTTCCCCTGCGCAACGACCACCC | 71 |
| A4 | ACGCGCCAGACGACAGTCGGTTGATGACACCCATTTTAGACCCTAAATAGCGGGCGGCGCAACGACCACCC | 71 |
| A5 | ACGCGCCAGACGACAGTTATGTGCCTTATCCAAAGCCGGGTGTAAAATCGCGGACCGCGCAACGACCACCC | 71 |

| | | |
|---|---|---|
| A6 | ACGCGCCAGACGACAGCATGGCACCTAAAAGGTGGCTTGTTGAGTACCCACGTTACGCGCAACGACCACCC | 71 |
| A7 | ACGCGCCAGACGACAGCTATAAGCTATGGCGGTTCACAGCTTCTGAGAACGTCGACGCGCAACGACCACCC | 71 |
| A8 | ACGCGCCAGACGACAGAGGTTGCACCATGGAGTTCCTTTCGTCACACAAGGACGTAGCGCAACGACCACCC | 71 |
| A9 | ACGCGCCAGACGACAGTTGAGATCGAAGTTCCCCGGATCAGTCGCTCATGAAGCTAGCGCAACGACCACCC | 71 |
| A10 | ACGCGCCAGACGACAGCCCAGCTCCGTTCATGGCATCAATAAAGGTTAGTGCGTGAGCGCAACGACCACCC | 71 |
| A11 | ACGCGCCAGACGACAGTTGGGTCGAAAACATGTGTGGTCTTCGCCTCCAAAGACCAGCGCAACGACCACCC | 71 |
| A12 | ACGCGCCAGACGACAGTTCAACGCTCAGTTGCGGTTCTGCAGCCAGTTGAAGAACAGCGCAACGACCACCC | 71 |
| **ssDNA for aptamers** | | |
| HA-LNA-1 | TCGGGCGTGTCGTCTGGCCTTTTGGCTTTGCTTTTCCCGCAATTCATCGTTTCCGTCCGCTTCCTCCTCCC | 71 |
| HA-LNA-2 | TCGGGCGTGTCGTCTGTCCTTTTGGCGTATCATTTTCATGTACATATCGTTTACGTCCGCTTCCTCCTCCC | 71 |
| HA-LNA-3 | TCGGGCGTGTCGTCTGCCCTTTTGTCAATCATCATTAATAATTGTATCGTTTGCGTCCGCTTCCTCCTCCC | 71 |
| hCD40L-1 | TCGGGCGTGTCGTCTGTGTGTCACGTCGTTTGACGTCCTTTTCACACCTGCTTTTTCCGCTTCCTCCTCCC | 71 |
| hCD40L-3 | TCGGGCGTGTCGTCTGTGTGCCCGCGCTTTCTTCTCTTTTTGGGAATCCCTGTTCCCCGCTTCCTCCTCCC | 71 |
| hCD40L-4 | TCGGGCGTGTCGTCTGTTGTTAATTCGTTTAAATTCCTTTTCAACTTGTTCATCGTCCGCTTCCTCCTCCC | 71 |
| hCD40L-6 | TCGGGCGTGTCGTCTGTCGTGAATCCCTGTTCACTTTGCCCTATTTTGTACGTTTTCCGCTTCCTCCTCCC | 71 |
